# Distant lymph nodes compensate for resected tumor-draining lymph nodes during cancer immunotherapy

**DOI:** 10.1101/2023.09.19.558262

**Authors:** Lutz Menzel, Hengbo Zhou, James W. Baish, Meghan J. O’Melia, Laurel B. Darragh, Derek N. Effiom, Emma Specht, Juliane Czapla, Pin-ji Lei, Johanna J. Rajotte, Lingshan Liu, Mohammad R. Nikmaneshi, Mohammad S. Razavi, Matthew G. Vander Heiden, Jessalyn M. Ubellacker, Lance L. Munn, Sana D. Karam, Genevieve M. Boland, Sonia Cohen, Timothy P. Padera

**Author notes:** Equal contribution authors (Lutz Menzel, Hengbo Zhou).

## Abstract

Lymphatic transport facilitates the presentation of cancer antigens in tumor-draining lymph nodes (tdLNs), leading to T cell activation and the generation of systemic anti-cancer immune surveillance. Surgical removal of tdLNs to control cancer progression is routine in clinical practice. However, whether removing tdLNs impairs immune checkpoint blockade (ICB) is still controversial. Our analysis demonstrates that melanoma patients remain responsive to PD-1 checkpoint blockade after regional LN dissection. We were able to recapitulate the persistent response to ICB after regional LN resection in murine melanoma and mammary carcinoma models. Mechanistically, soluble antigen is diverted to distant LNs after tdLN dissection. Consistently, robust ICB responses in patients with head and neck cancer after primary tumor and tdLN resection correlated with the presence of reactive LNs in distant sites. These findings indicate that distant LNs sufficiently compensate for the removal of direct tdLNs and sustain the response to ICB.

## Introduction

Advances in our understanding of the mechanisms of T cell activation and exhaustion led to the identification of several immune checkpoints, such as programmed death 1 (PD-1)^1,2^ and cytotoxic T lymphocyte antigen 4 (CTLA-4)^3,4^, which are now therapeutic targets^5,6^. Immune checkpoint blockade (ICB) has been approved for the treatment of many cancers, including melanoma^7,8^ and breast cancer^9,10^. However, an improved understanding of the mechanisms of ICB therapy is necessary to benefit more patients.

Lymph nodes (LNs) surveil tissue fluids and initiate adaptive immunity^11,12^. Lymphatic vessels collect interstitial fluid and cells to create lymph and transport the lymph to draining LNs. The tissue region that drains to a specific group of LNs is called a lymphosome. As lymphatic vessels can take up cells— including cancer cells—metastatic dissemination to tumor-draining lymph nodes (tdLNs) can occur during the early stage of tumor growth^13^. Colonization in LNs often impairs nodal entry of naïve lymphocytes^14^, causes expansion of regulatory T cells (Tregs)^15–17^ and induces systemic tolerance in favor of cancer progression^16^. LN metastases can also disseminate to distant metastatic sites^18,19^, further driving cancer progression.

Clinically, sentinel lymph node biopsy (SLNB) and completion lymph node dissection (CLND) are commonly performed^20–24^ to assess the disease stage and to limit cancer progression. Seminal work demonstrated that tumor-specific T cells and non-exhausted tumor-specific memory T cells are enriched in tumor-draining, but not in other LNs^25,26^, suggesting that removal of tdLNs would impair anti-cancer immune responses. However, in a phase III clinical trial, melanoma patients treated with tumor, sentinel LN and metastatic LN resection still responded to anti-CTLA4 therapy, indicating that the presence of tdLNs may not be necessary to maintain ICB efficacy^27^. Similarly, after SLNB, administering anti-CTLA4 locally to the lymphatic basin at the site of the resected tumor enhanced effector T cell activation, lasting up to 3 months. These data suggest that the ICB-mediated immune response extends beyond just the sentinel LNs^28^. Due to the important role of LNs in the adaptive immune response, it is imperative to evaluate the effects of SLNB and CLND on ICB efficacy. Our retrospective analysis demonstrates that stage III melanoma patients that underwent CLND still benefit from adjuvant anti-PD1 inhibition. Using orthotopic breast cancer and melanoma models in mice, we were able to show that ICB efficacy persists after tdLN dissection as a result of a compensatory mechanism by which tumor-derived antigen is diverted to distant LNs (non-tdLNs). This phenomenon was corroborated in head and neck cancer patients who underwent neoadjuvant ICB therapy followed by bilateral neck dissection, radiation and adjuvant ICB, where the ICB response was associated with the presence of reactive LNs at distant sites. To apply our findings to improve treatment strategies, we compared systemic versus locoregional administration of ICB and found that delivering ICB to the lymphosome of reactive non-tdLNs improves anti-tumor T cell responses after CLND.

## Results

### Stage III patients with melanoma respond to ICB after CLND

Prior to the availability of effective systemic therapies for patients with melanoma, LN surgery played a major role in the treatment of patients with locoregional disease. Any patient with clinically evident LN metastases or a positive SLNB underwent CLND for disease control. In 2017, MSLT-II^29^ (a randomized clinical trial of CLND vs observation in early-stage melanoma patients with a positive SLNB) demonstrated that CLND did not improve melanoma-specific survival (MSS). Due to the morbidity associated with CLND, this trial led to a change in clinical practice, favoring observation for early-stage patients with melanoma with a positive SLNB. CLND is now reserved for cases of disease recurrence in the draining LN basin. Approximately 20% of patients will eventually experience recurrence in the draining nodal basin, while 80% of patients with a positive SLNB are thus spared the unnecessary morbidity risk of CLND^29^. Given the importance of tdLNs for ICB response in mouse models^26,30,31^ and humans^32^, we asked whether the extent of LN surgery impacts the effectiveness of ICB treatment. Our retrospective analysis included 66 patients with stage III melanoma treated with surgery and adjuvant anti-PD1 therapy (nivolumab or pembrolizumab), from which 16 underwent CLND and 50 underwent only SLNB (**Figure S1A**). Surprisingly, there was no statistically significant difference in melanoma specific survival (MSS) (**Fig. 1A**) or recurrence-free survival (RFS) (**Fig. 1B**) between both groups. Furthermore, no significant differences were observed after stratification by local recurrence-free survival (**Fig. 1C**) and distant recurrence-free survival (**Fig. 1D**). To validate these findings in patients with metastatic disease in the tdLN basin, we narrowed our cohort to patients with a positive SLNB (**Figure S1A**). Of note, the number of patients undergoing CLND is minimal (**Figure S1B**) as a result of the MSLT-II trial findings—which demonstrated that CLND provided no (MSS) benefit^29^. We selected stage IIIC patients to assess whether the extent of nodal surgery in patients with known metastatic melanoma in the draining nodal basin affected the response to anti-PD1 therapy (**Figure S1C-E**). Anti-PD1 therapy significantly enhanced the RFS in patients who did not undergo lymph node surgery, consistent with clinical trial data (**Fig. 1E)**. However, LN dissection did not impair the efficacy of anti-PD1 treatment (**Fig. 1E**). Our data are limited to our specific patient population, and it is important to note our study was not designed to evaluate whether CLND has a therapeutic benefit. However, our findings were confirmed in a recently reported retrospective analysis of sentinel LN positive, stage III melanoma patients treated with adjuvant anti-PD-1 monotherapy that underwent CLND or no LN dissection. Multivariate analysis of 261 patients from 39 independent institutions receiving adjuvant anti-PD-1 treatment showed no difference in 2-year overall survival (OS) or RFS with or without CLND (2-year RFS: 55.9% vs. 52.6%, p=0.74; 2-year OS: 83.4% vs. 83.6%, p=0.73)^33^. These data demonstrate that the response to ICB persists in patients after SLNB and CLND surgery.

**Figure 1.**
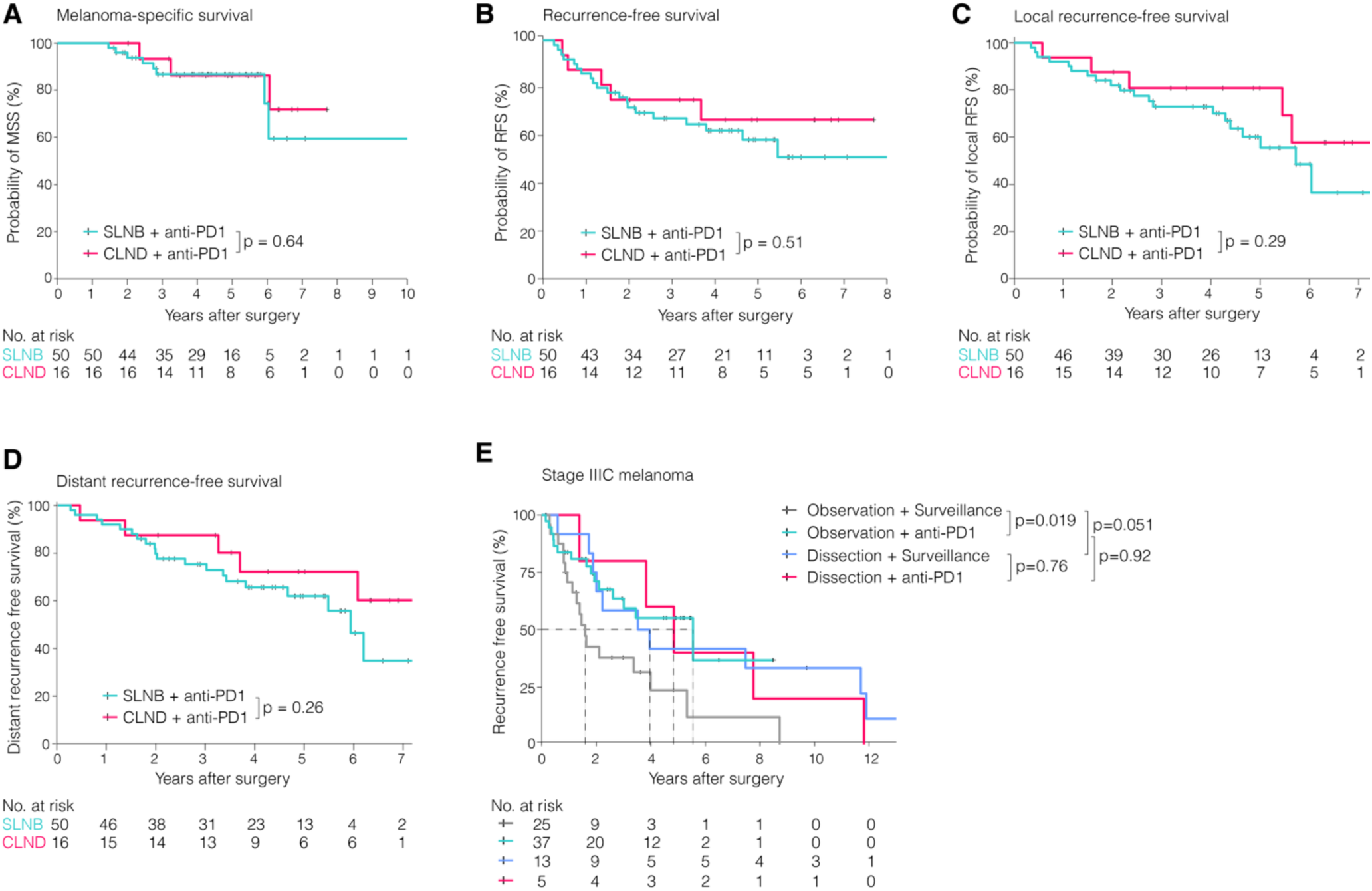
Adjuvant ICB response persists in stage III melanoma patients after LN surgery. Retrospective analysis with a cohort of cutaneous stage III melanoma patients who underwent SLNB or CLND, treated with adjuvant anti-PD1 antibody (nivolumab or pembrolizumab) at the Massachusetts General Hospital. **(A)** Melanoma-specific survival, **(B)** recurrence-free survival, **(C)** local recurrence-free survival and **(D)** distant recurrence-free survival. **(E)** Retrospective analysis of stage IIIC melanoma patients with observation or LN dissection and either surveillance or anti-PD1 treatment. Statistical analysis with Log-rank test. See also Figure S1.

### ICB response persists after tdLN dissection in mice

To determine if the sentinel LNs or a combination of the sentinel and adjacent LNs are required for efficacy of ICB response, we orthotopically implanted murine melanoma cells (intradermal injection of B16F10 or YUMMER1.7) or murine breast cancer cells (mammary fat pad injection of E0771). We followed a treatment regimen similar to that described in recent studies^26,30^, in which mice received three doses of either IgG or anti– PD-1 every two to three days, starting on day 7 after tumor implantation. Using this schedule, we found that micrometastases could still be detected in the tumor-draining lymph node (tdLN) following ICB therapy. These findings highlight the potential for residual metastatic cells to persist in lymphatic tissues despite ICB treatment (**Figure S2A-C)**. To test whether removal of tdLNs alters ICB responses, we performed sham surgeries, surgically removed the ipsilateral inguinal (SLNB) or a combination of the ipsilateral inguinal (inLN), axillary (axLN), and brachial (brLN) LNs (CLND) 6 days after tumor implantation. Subsequent treatment with anti-PD1 antibody resulted in a significant tumor growth delay for E0771 and YUMMER1.7 tumors (**Figure S2D**) and increased the survival in all three tumor models compared to IgG controls (**Fig. 2A**).

**Figure 2.**
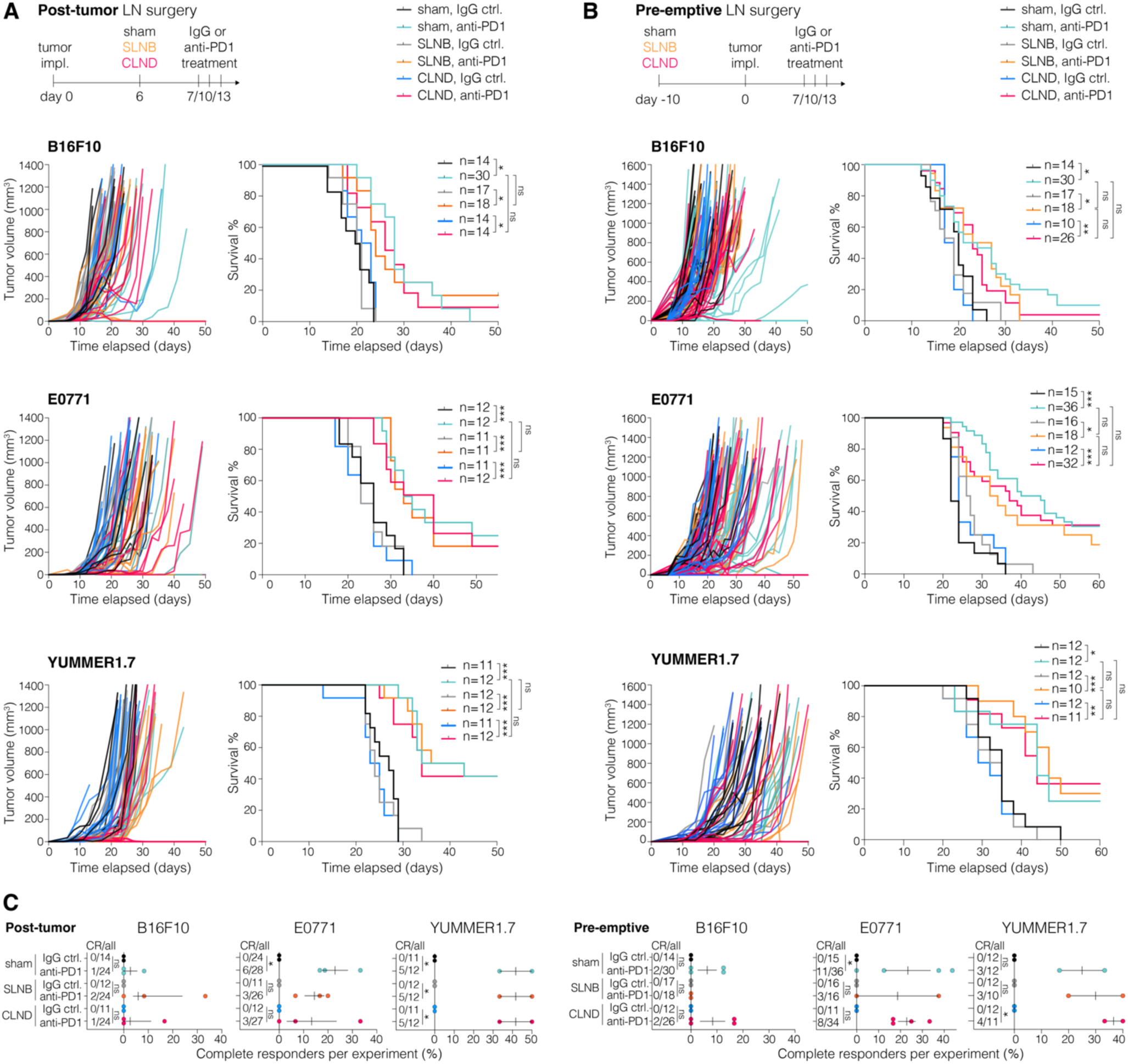
ICB response persists after LN dissection. **(A)** Orthotopically implanted B16F10, E0771 or YUMMER1.7 tumors in mice with sham surgery, SLNB, or CLND and IgG control or anti-PD1 treatment. **(B)** Pre-emptive sham surgery, SLNB, or CLND in mice with orthotopic B16F10, E0771 or YUMMER1.7 tumor implantation and IgG control or anti-PD1 treatment. **(C)** Data points represent the average percentage of complete responders (CR) per experiment. CR as a fraction of the total number of mice are indicated as CR/all. Number of mice per group indicated as n from 2-3 individual experiments. Statistical analysis with Log-rank test (A,B) or Fisher’s exact test (C), ns=not significant \**p*<0.05, \*\**p*<0.01, \*\*\**p*<0.005. See also Figure S2.

To determine whether 6 days of priming the tdLNs was sufficient to generate systemic anti-cancer immunity, we used pre-emptive LN removal similar to that used in recent studies^26,30^. We performed SLNB or CLND before tumor implantation and again found that LN resection had no negative effect on ICB response (**Fig. 2B**). The frequency of complete responders (CR) with anti-PD1 treatment was similar to IgG controls in the weakly-immunogenic B16F10 tumor model but showed significant differences in the E0771 and YUMMER1.7 tumor models (**Fig. 2C**). The extent of LN surgery did not alter the frequency of CRs in anti-PD-1 treated animals in any of the tumor models tested. (**Fig. 2C**). These findings are not consistent with recent studies using MC38 and CT26 tumor models grown ectopically in a subcutaneous setting^26,30^. To evaluate this discrepancy, we implanted E0771 ectopically in the dermis and showed that CLND was able to reduce the response to anti-PD1 (**Figure S2E,F**), consistent with other studies of ectopic tumors. These data indicate that altered immune response can occur when tumor cells are implanted ectopically^34,35^.

The lack of an effect of CLND when tumors are grown orthotopically prompted us to examine whether LNs are even necessary for ICB efficacy or if other lymphoid organs like the spleen compensate for resected LNs. Experiments including splenectomy in combination with CLND showed that the spleen was not significantly involved in the sustained ICB responses after LN dissection (**Figure S2G**). In contrast, responses to ICB treatment were impaired in *Lta^−/−^* mice, which do not develop peripheral LNs, while maintaining normal circulating lymphocyte populations^36–38^ (**Figure S2H**). These data suggest that effective ICB treatment requires LNs in general, but not necessarily the direct tdLNs.

### Removal of draining LNs promotes antigen transport to distant LNs

As LNs are critical for the presentation of antigen by dendritic cells to naïve T cells, we hypothesized that antigen might be transported to alternative LNs after CLND. To interrogate this hypothesis as a compensatory mechanism, we injected surrogate antigens (fluorophore-conjugated dextran of 4.4 kDa or 2,000 kDa) into the 4^th^ mammary fat pad (MFP) to measure the local and systemic transport to alternative LNs after SLNB or CLND. Most of the 2,000 kDa dextran aggregated in the LN subcapsular sinus (SCS) (**Fig. 3A**) and appeared within lymphatic vessels of the LN medulla (**Fig. 3B**) confirming that molecules of this size are predominantly transported via lymph flow. 4.4 kDa dextran entered the LN parenchyma and was distributed throughout the paracortex (**Fig. 3A**), consistent with entry via blood. Small biomolecules are known to access the blood circulation, whereas molecules larger than 40 kDa transport more through lymph and are retained within the SCS when they arrive in the LN^39^. Six hours after injection, dextran levels in axillary LNs following SLNB matched those in sham inLNs, with negligible tracer in contralateral nodes. In contrast, CLND led to detectable dextran in the contralateral inLN at six hours (**Fig. 3C**), demonstrating its diversion to distant LNs.

**Figure 3.**
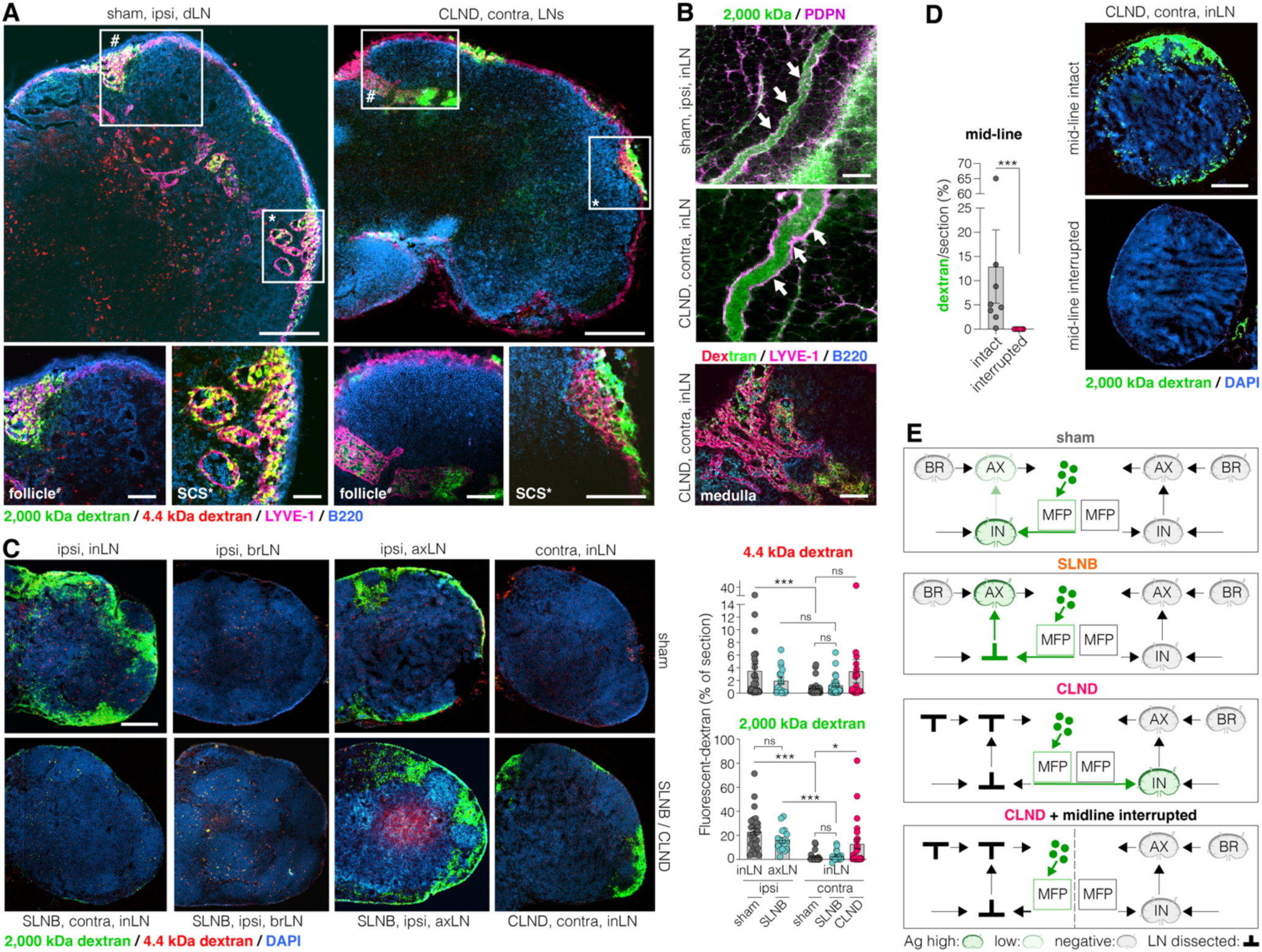
Dissection of draining LNs promotes antigen transport to distant LNs. **(A)** Representative images demonstrate intranodal dextran distribution on the ipsilateral side after sham surgery or contralateral side after CLND. Subcapsular sinus (SCS). Scale bar: 250 µm (top row), 100 µm (bottom row). **(B)** *Top*, Representative images of 2,000 kDa FITC-dextran within lymphatic vessels (Podoplanin, PDPN) and, *bottom*, 2,000 kDa FITC-dextran and 4.4 kDa TRITC-dextran in the LN medulla (LYVE-1) in contralateral LNs after CLND. Scale bar: 20 µm (*top*), 200 µm (*bottom*). **(C)** 2,000 kDa FITC-dextran and 4.4 kDa TRITC-dextran were injected into the mammary fat pad (MFP) after sham surgery, SLNB or CLND. Dextran distribution 6 hours post-injection in axillary (axLN), brachial (brLN) and inguinal (inLN) LNs after sham, SLNB or CLND. Quantification of the fluorescent area (FITC or TRITC) in percent of whole section. Scale bar: 200 µm. **(D)** 2,000 kDa FITC-dextran in contralateral inLN 6 hours post-injection into the ipsilateral MFP of mice after CLND with intact or interrupted (cut) ventral skin at the midline. Scale bar: 200 µm. **(E)** Illustration of lymph flow distribution patterns after different surgery settings. Graphs show mean ± SEM with data points representing individual animals. Statistical analysis with Mann-Whitney U test (C,D), ns= not significant, \**p*<0.05, \*\*\**p*<0.005. See also Figure S3.

To further test whether antigen access to distant LNs can be enhanced when local lymphatic drainage is ablated, we dissected the popliteal LN (poLN), inLN, axLN and brachial LN (brLN) of the ipsilateral side (ILND) and injected the dextran into the ipsilateral foot pad. By avoiding injection-induced lateral flow of antigen by food pad injection, we were able to monitor compensatory lymph drainage after ILND. Similar to results in **Fig. 3C**, we discovered that the 2,000 kDa dextran was mainly located within the ipsilateral poLN, inLN and axLNs when lymphatic drainage was intact (sham), whereas ILND diverted the lymph transport to the contralateral inLN and axLN (**Figure S3A,B**). The absence of 2,000 kDa dextran in the contralateral poLNs suggests that, once taken up by lymphatic vessels of an adjacent lymphosome, lymph-borne antigens follow normal drainage patterns and do not undergo retrograde flow or circulate significantly through the blood. In contrast, 4.4 kDa dextran was found in all the remaining LNs (**Figure S3A, B**), suggesting transport through lymph and blood that results in a systemic distribution. The compensatory drainage phenomena suggested by our data have also been reported clinically, including evidence that lower limb interstitial fluid can cross the lower abdomen towards the contralateral inguinal region^40–42^. Corresponding to these reports, we next tested if the interstitial transport from a disrupted lymphosome crosses the midline and reaches contralateral LNs using a dermal route. To do so, we surgically cut the skin along the midline (lower abdomen to chest) in CLND mice to interrupt intradermal transport and then injected dextran into the ipsilateral MFP. Dextran transport to contralateral LNs was detected in mice with intact midline skin but was absent after midline incision **(Fig. 3D)**. These data indicate that interstitial fluid can move across the midline through the dermis and reach an adjacent lymphosome draining to distant LNs after CLND. In summary, after removal of the primary dLNs antigen moves through interstitial tissues to the edge of an adjacent lymphosome with intact lymphatic drainage, where the antigen is taken into initial lymphatic vessels and follows the normal lymphatic drainage to the associated LN. Through this dermal transport mechanism, antigen is able to cross the midline to intact lymphatics that drain to the contralateral LNs after CLND (**Fig. 3E**).

### Mathematical modeling shows antigen access to adjacent lymphosomes via dermal interstitial transport after tdLN dissection

Our data show that interstitial antigen enters adjacent lymphosomes after dissection of one or more LNs (**Fig. 3**). On this basis, we developed anatomically-based, computational models of interstitial fluid pressure (IFP) and lymph flow rates that predict how antigens and large tracer molecules can be redistributed to alternative LNs when superficial lymphosomes are disabled by removing their draining lymph nodes. The models recapitulate our primary experimental findings in mice (**Fig. 4A-F**) and predict how these results may translate to clinical practice in humans (**Fig. 4G-I**). Details of the model are provided in **Supplemental Materials** along with analytical validation and parametric studies (**Figure S4**)

**Figure 4.**
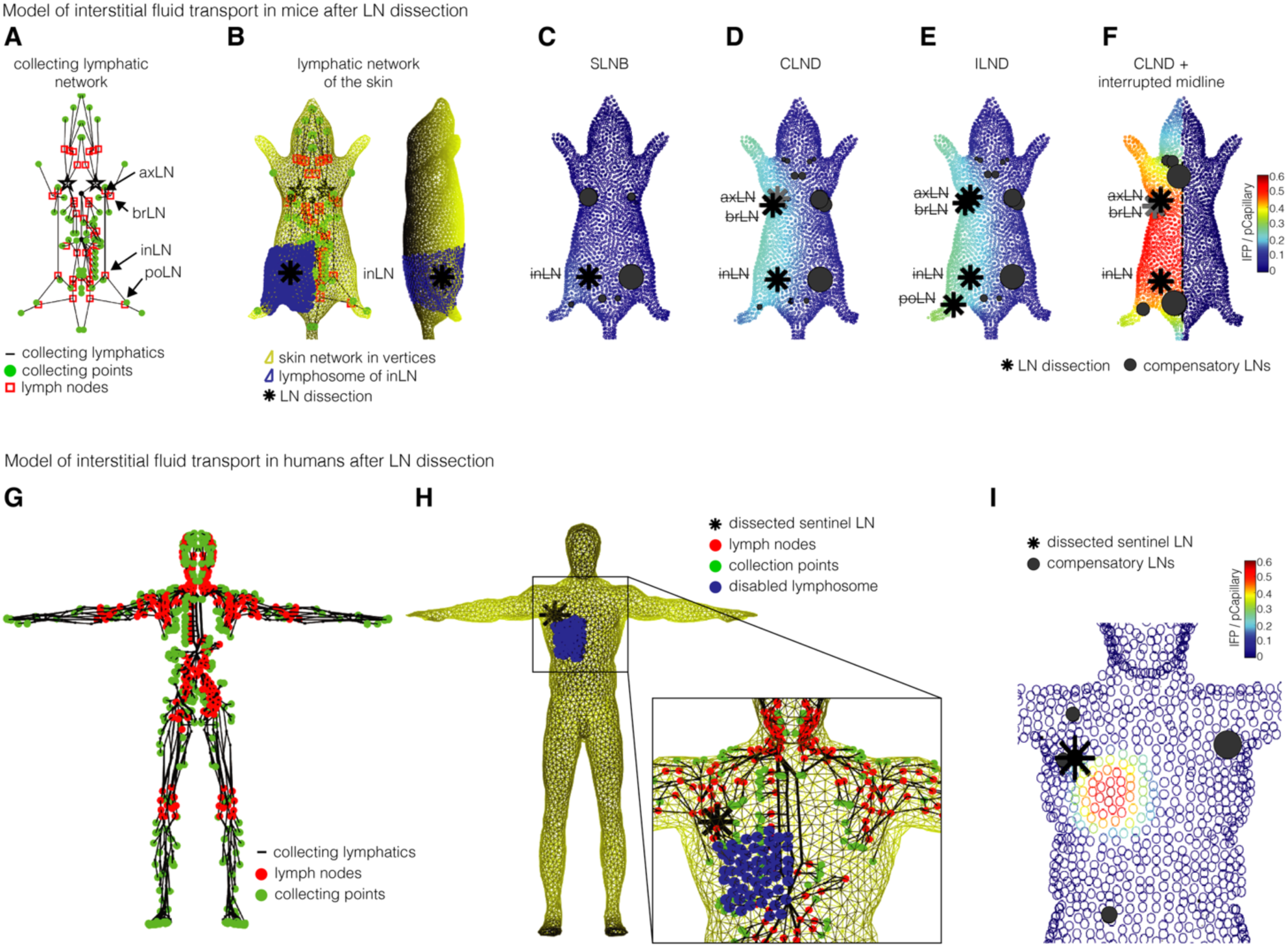
Mathematical modeling of antigen access to adjacent lymphosomes via dermal transport after node dissection. **(A)** Mouse model with the network of collecting lymphatic vessels (black), collecting points (blue) and LNs (red). **(B)** Computational mesh (yellow) for the network of initial lymphatic vessels in the skin showing the drainage basin (lymphosome) (blue) for the right inguinal node (*) **(C-F)** Color maps of the interstitial pressure relative to the capillary blood pressure for disabled nodes (*) corresponding to **(C)** SLNB (inLN), (**D**) CLND (in/ax/br), **(E)** ILND (po/in/ax/br), **(F)** CLND (in/ax/br) with severed midline. The dark circles represent compensatory lymph nodes and are scaled by the fraction of diverted flow they must carry. **(G)** Human male model with the network of collecting lymphatic vessels (black), collecting points (green) and LNs (red). **(H)** Computational mesh (yellow) for the network of initial lymphatic vessels in the skin showing the drainage basin lymphosome (blue) for a right axillary node (*). **(I)** Color map of the interstitial pressure relative to the capillary blood pressure for a disabled right axillary node (*) with dark circles representing the compensatory LNs scaled by the change in flow induced by the diversion of flow from the disabled lymphosome. See also Figure S4.

The color maps in **Fig. 4C-F, I** show how disabling various combinations of the inLN, axLN, brLN and poLN causes local increases in interstitial fluid pressure that divert interstitial fluid to adjacent lymphosomes where it can be absorbed by active lymphatic vessels. The size of the black circles in **Fig. 4C-F, I** indicates the fractional increase in flow to other lymph nodes when selected lymph nodes were disabled. Normally with an intact lymphatic system, an equilibrium exists in which fluid lost from the blood capillaries is absorbed by a highly-connected network of valve-less, noncontracting initial lymphatic vessels and then actively pumped to local lymph nodes via collecting lymphatic vessels. Under normal conditions, interstitial fluid pressures remain at or slightly below atmospheric pressure with little lateral movement of fluid occurring in the superficial tissue layers. In contrast, when local lymph nodes were computationally removed, accumulating fluid increases the local interstitial pressure inducing passive flow through the dermal initial lymphatic vessels and dermis from regions of high interstitial pressure to distal regions with normal, low interstitial pressure and intact draining LNs. Typically, disabling a lymph node changed the flow most significantly in adjacent lymphosomes, although flow was also perturbed farther away. **Fig. 4F** shows that numerically severing the skin flow along the midline prevents flow changes on the contralateral side as we observed experimentally (**Fig. 3D**).

Our results in a human model (**Fig. 4G-I**) were qualitatively similar to those in mice, but important differences arise because of the relative size of the lymphosomes, anatomical connectivity of the vessels and the degree of redundancy in human lymphatic networks. When compared to the large lymphosomes found in humans, the small lymphosomes found in mice drain smaller quantities of fluid over shorter distances and generate only modest increases interstitial pressure when a LN is disabled. For example, the low IFP (light blue) in **Fig. 4C** for a disabled inguinal LN in a mouse can be compared to the much higher IFP (red) at the center of the human chest when axillary drainage is disabled in **Fig. 4I**. This size difference alone may account for much of the relative difficulty in creating experimental models of lymphedema in laboratory mice.

To further examine the effect of body composition and anatomical size, we compared the IFP in the skin from the human 3D model with an analytical assumption of a planar, circular lymphosome with equal surface area (**Figure S4**). Parameters, such as blood pressure, capillary permeability, dermal permeability, and lymphatic pumping capacity—which are affected by obesity, fitness and physical activity—can alter the interstitial pressure and volume of diverted flow (**Figure S4A-C)**. We also found that the human 3D model and the analytical assumption of a planar, circular lymphosome with equal surface area resulted in similar distribution patterns of IFP relative to lymphosome size (**Figure S4D**). To examine the size dependence of lymphosomes, we performed a parametric study of the model and found that larger lymphosomes, such as those found in humans, collect a higher volume of interstitial fluid resulting in higher IFP, which then maintains higher lymph flow (**Figure S4E**). The model demonstrates how the mobility of the fluid (***κ***) in the superficial network of initial lymphatic vessels affects the IFP but has little influence on the total quantity of antigen-carrying fluid diverted elsewhere from its initial destination **(Figure S4F)**. The similar lymph fluid diversion patterns and transport to compensatory LNs between the experimental mouse model and the mathematical model of human antigen drainage suggest our results are relevant for predicting lymph drainage in patients with or without LN dissection. The greater number of lymph nodes in the axillary and inguinal regions in humans compared to mice also alters the likely effects of removing any single lymph node.

### Antigen diversion to distant LNs promotes DC–driven immune responses after CLND

Having established that interstitial antigens drain to contralateral LNs following CLND, we next examined the role of dendritic cells (DCs) in antigen presentation. B16F10^OVA-ZsGreen^ and E0771^OVA-ZsGreen^ tumors implanted in mice treated with LN surgery and ICB (**Figure S5A**) grew significantly slower in the anti-PD1 group with no detrimental effect of CLND (**Fig. 5A**). To assess DC uptake of tumor-derived antigen, we employed tumor cell–expressed ZsGreen, which was found in all tumor cells and in the majority of intratumoral DCs (**Fig. S5B**). In the melanoma model, the ipsilateral inLN after sham surgery had the highest proportion of zsGreen^+^ DCs, while the ipsilateral axLN only contained zsGreen^+^ DCs after SLNB. The contralateral inLN only contained ZsGreen^+^ DCs after CLND in the melanoma model. Growing within the MFP, the breast cancer model caused ZsGreen^+^ DCs on the ipsilateral side within the inLN but also within the axLN, independent of SLNB. The location close to the body midline also resulted in some—but still low numbers—ZsGreen^+^ DCs in the contralateral inLN during sham and SLNB conditions. These data are in contrast to the results from melanoma grown in the more lateral flank skin. However, CLND still caused a significant increase of ZsGreen^+^ DCs in the contralateral inLN in the breast model (**Fig. 5B**). We confirmed these findings by staining ZsGreen^+^ DCs for ovalbumin antigen presented on MHC-I (MHC-I:SIINFEKL), showing significantly more SIINFEKL-presenting DCs within the ipsilateral LNs with sham surgery and contralateral LNs of mice with CLND (**Figure S5D**). DCs and macrophages (MΦ) in the spleen appeared to be mostly unaffected by the tumor model and surgery group (**Fig. 5B, Figure S5E**).

**Figure 5.**
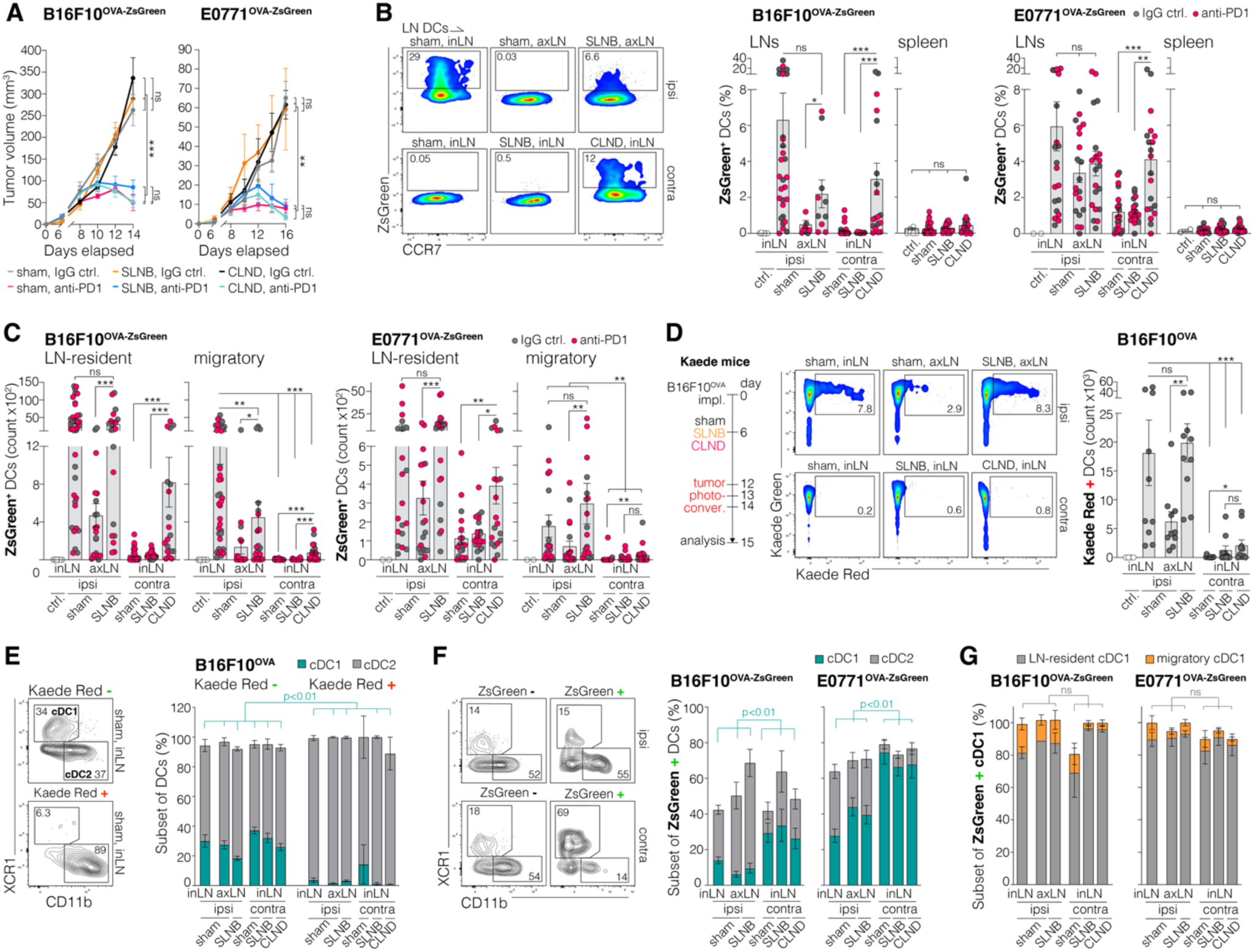
Antigen diversion to distant LNs promotes DC–driven immune responses after CLND. **(A)** Volume of B16F10^OVA-ZsGreen^ or E0771^OVA-ZsGreen^ tumors, n=11-12 per group in 2 individual experiments. Corresponding to flow analysis in B, C, F. Experimental set up and gating strategy illustrated in Figure S5A-C. **(B)** Representative flow cytometry plots and quantification of the number of ZsGreen^+^ DCs per LN. **(C)** Number of ZsGreen^+^ migratory and LN-resident DCs show LN-resident DCs are the dominant ZsGreen^+^ population in the contralateral LNs after CLND. **(D)** Quantification of Kaede Red^+^ DCs in LNs of Kaede mice with photoconverted B16F10^OVA^ tumors show limited migration of DCs from the primary tumor to contralateral LNs. Gating strategy in Figure S5C. **(E)** cDC1/cDC2 proportions of Kaede Red^+^ tumor-derived DCs and Kaede Red^−^ DCs. Statistics indicated for cDC1. **(F)** cDC1/cDC2 proportions of ZsGreen^+^ DCs in LNs of mice with B16F10^OVA-ZsGreen^ or E0771^OVA-ZsGreen^ tumors, including IgG control and anti-PD1 treated animals (n=10-30/group). Statistics indicated for cDC1s. **(G)** LN resident and migratory proportions of ZsGreen^+^ cDC1s, as in (F), in LNs of mice with B16F10^OVA-ZsGreen^ or E0771^OVA-ZsGreen^ tumors, including IgG control and anti-PD1 treated animals (n=6-35/group). Statistics indicated for LN-resident cDC1. Graphs show mean ± SEM, data points representing individual animals. Statistical analysis with Mann-Whitney U test (B, C, D), t-test (E,F,G) or 2-way ANOVA (A), ns= not significant, \**p*<0.05, \*\**p*<0.01, \*\*\**p*<0.005. See also Figure S5.

We further investigated whether tumor antigen presenting DCs in compensatory LNs were primarily LN-resident DCs (resDC) that took up soluble antigens or migratory DCs (migDC, CD103^+^/CCR7^+^) that carried the tumor antigen from the primary tumor. Our results indicate that migDCs comprise a relevant proportion of tumor antigen carrying DCs in the sentinel LNs, inLN for the melanoma model and inLN/axLN for breast cancer. CLND increased the numbers of ZsGreen^+^ migDCs slightly—though they were still many fewer than ZsGreen^+^ resDCs—in the contralateral inLN (resDC 10x more than migDC) (**Fig. 5C**). Photoconversion of B16F10^OVA^ melanoma tumors in Kaede mice **(Fig. S5C,F-G)** confirmed our findings that even though CLND causes a slight increase in tumor-derived migDCs in contralateral inLNs, migDCs were almost exclusively found in LNs on the ipsilateral side, primarily within the sentinel inLN or the axLN after SLNB (**Fig. 5D**). These data are consistent with our antigen transport data and suggest that soluble antigen reaches the contralateral LNs after CLND where it is presented by LN resident DCs. The contribution of migratory DCs to contralateral antigen presentation is minimal.

Because cDC1s (CD11b^−^/XCR1^+^)^43^ almost uniquely cross-present phagocytosed tumor antigens, they are the main initiators of antitumor CD8^+^ T cell responses^44^. In the Kaede Red negative (non–tumor-derived) DC pool, we saw a stable cDC1:cDC2 ratio of ∼30:60 across all LNs and conditions, whereas tumor-derived Kaede Red^+^ DCs were almost exclusively cDC2s across all LNs and conditions (**Fig. 5E**). In WT mice bearing B16F10^OVA-ZsGreen^ or E0771^OVA-ZsGreen^ tumors, ZsGreen^+^ antigen-carrying cells on the contralateral side showed a marked enrichment of cDC1s (**Fig. 5F**), nearly all of which were LN-resident (**Fig. 5G**).

Our data indicate that interstitial antigens reach the contralateral LNs after CLND and are mostly presented by LN-resident cDC1s, with only few antigen-carrying migratory DCs contributing. This is in contrast to the ipsilateral sentinel LN, where resident and migratory DCs are equally involved in presenting tumor antigens.

### Locoregional ICB administration enhances T-cell responses in distant LNs

We next examined whether T cell responses are generated in distant LNs after CLND. After B16F10^OVA-zsGreen^ or EO771^OVA-zsGreen^ implantation, ipsilateral LNs from sham surgery mice and contralateral LNs from CLND mice were significantly expanded in volume and in CD45^+^ leucocytes, while the proportional immune cell composition remained similar between groups (**Figure S6A**). Concomitantly, the number of antigen-experienced (CD44^+^PD1^+^) T cells was significantly higher in ipsilateral LNs from sham surgery mice and contralateral LNs from CLND mice with the numbers further increasing with anti-PD1 treatment after CLND (**Fig. 6A**). Confirming the latter, OVA-tetramer^+^ T cells (OVA-T cells) were enriched during B16F10^OVA-zsGreen^ tumor growth in ipsilateral LNs of sham mice and contralateral LNs of CLND mice but almost absent in contralateral or cervical LNs of sham mice (**Fig. 6B**), emphasizing that CLND induces regional redistribution of antigen but not systemic antigen diversion. The abundance of OVA-T cells was similar in spleens and blood of sham and CLND mice after anti-PD1 treatment (**Fig. S6B**), demonstrating that the generation of circulating tumor-specific T cells persists after LN surgery.

**Figure 6.**
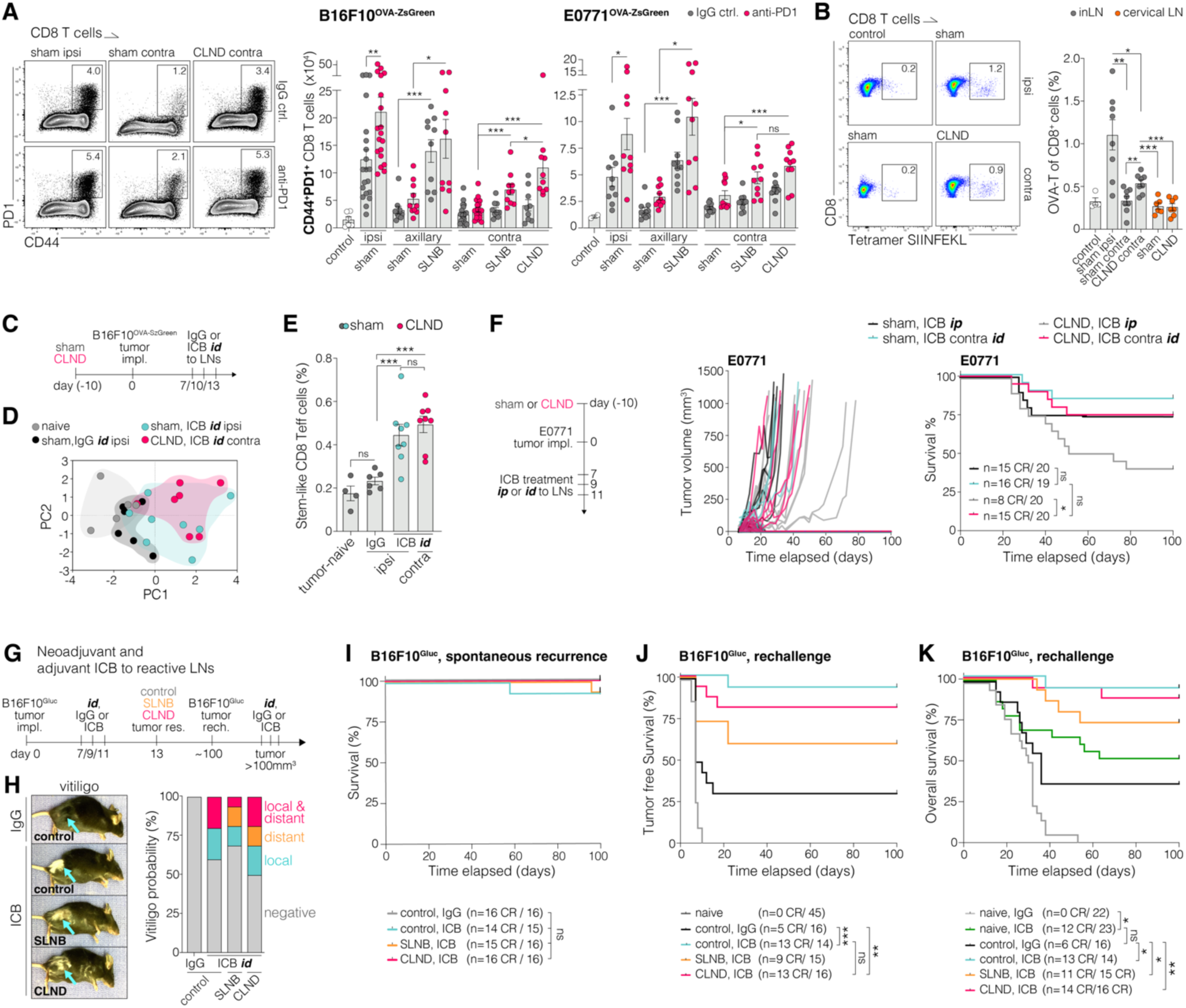
Administration of ICB to distant LNs improves response after CLND. **(A)** Representative flow cytometry plots and number of antigen-experienced (CD44^+^PD1^+^) CD8 T cells per LN from mice with B16F10^OVA-ZsGreen^ or E0771^OVA-ZsGreen^ tumors. Controls represent mice without tumor implantation. **(B)** Tetramer^+^ (MHCI-SIINFEKL) CD8 cells in LNs of mice with B16F10^OVA-ZsGreen^ tumors. Controls represent mice without tumor implantation. **(C)** Experimental set up, pre-emptive sham surgery or CLND before orthotopic E0771 tumor implantation. IgG control or a combination of anti-PD1 and anti-CTLA4 were intradermally (*id*) administered to the ipsilateral LNs or contralateral LNs (referred to as ICB). Corresponding to D,E. **(D)** Principal component analysis of T cells from respective LNs. Analysis based on the abundance of antigen-experienced T cells (CD44^+^, PD1^+^), stem-like T cells (CD44^+^, PD1^+^, TCF1^+^), central memory T cells (CD44^+^, CD62L^+^), effector memory T cells (CD44^+^, CD62L^−^, PD1^−^), and stem-like effector memory T cells (CD44^+^, PD1^+^, TCF1^+^, CD62L^−^, CD39^−^). **(E)** Percentage of stem-like effector memory T cells in all T cells. **(F)** Tumor volume and overall survival of mice with E0771 tumors. Neo-adjuvant ICB was administered *ip* or *id* to the contralateral inLN. n indicates the number of mice per group of at least 2 experiments, complete responders (CR). **(G)** Experimental set up of neoadjuvant-adjuvant treatment of B16F10^Gluc^ and tumor rechallenge. Control group represents tumor resection without LN surgery. Corresponding to I,J,K. **(H)** Representative images of vitiligo (arrows) developed in mice after neoadjuvant treatment and quantification of mice with local, distant or local and distant vitiligo relative to the side of tumor implantation. **(I)** Overall survival during spontaneous tumor recurrence. **(J)** Tumor-free survival of mice previously treated with neoadjuvant ICB, primary tumor resection and tumor rechallenge. Naive and untreated mice were used as controls for tumor rechallenge. **(K)** Overall survival of mice treated with neoadjuvant-adjuvant ICB after the rechallenge. Number of mice per group indicated as n from 2-3 individual experiments. Graphs show mean ± SEM with data points representing individual animals. Statistical analysis with Mann-Whitney U test (A,B,E) or Log-rank test (F,I,J,K), ns= not significant, \**p*<0.05, \*\**p*<0.01, \*\*\**p*<0.005. See also Figure S6 and Figure S7.

We further tested the T cell response in an adjuvant setting. We resected primary tumors in combination with sham or CLND surgery, followed by subsequent anti-PD-1 treatment. There was no difference in abundance of OVA-T cells between ipsilateral LNs from sham surgery mice and contralateral LNs from CLND mice (**Fig. S6C**), indicating that distant LNs can generate tumor-specific T cell responses even after resection of the primary tumor. In addition to T cell responses in LNs, we analyzed primary tumor-infiltrating T cells and found that anti-PD1 treatment enhanced proliferating effector T cells in sham and CLND mice (**Figure S6D,E,H**). Adjuvant anti-PD1 treatment of sham and CLND mice induced the generation of a memory T cell population upon tumor rechallenge. Non-exhausted stem-like central memory CD8 T cells were similarly abundant in ipsilateral LNs of sham mice and contralateral LNs of CLND mice (**Figure S6F,I**). Using photoconversion of the contralateral LN in Kaede mice after rechallenge with B16F10^OVA-zsGreen^ tumors, we confirmed that tumor-specific T cells from distant LNs traffic to the rechallenge tumors after CLND (**Fig. S6G**). Taken together, ICB-mediated expansion of tumor-infiltrating T effector and T memory cells persist after CLND.

Previous studies demonstrated that locoregional ICB augments cancer immunotherapy^28,45,46^. These findings, together with our results, emphasize the potential of distant LNs as ICB target sites. Because presentation of tumor antigens in distant LNs was almost only present after dissection of the primary tdLNs, we assessed the capacity of contralateral LNs to respond to locoregional ICB injection in mice after CLND (**Fig. 6C**). As anti-PD1 as a single target is based on a mechanism primarily effective within tumor tissue^47^, we used the clinically effective combination of anti-PD1 and anti-CTLA4 treatment^48^, which is more likely to have an effect during APC–T cell encounter in LNs^47^. Intradermal (*id*) administration of ICB in the lymphosome drained by the ipsilateral tdLNs with sham surgery or administration in the contralateral lymphosome in mice with CLND resulted in robust LN expansion in both groups after ICB but not in IgG treated controls (**Figure S7A**). Within the T cell compartment, ipsilateral LNs of sham mice and contralateral LNs of CLND mice shared a similar composition of T cell subsets based on principal component analysis (**Fig. 6D**), including a similar abundance of stem-like effector T cells, which are considered to be the main contributors to effective ICB responses^49^ (**Fig. 6E**). To further investigate the potency of distant LNs for ICB treatment, we performed sham surgery or CLND in mice with E0771 tumors followed by ICB administration either intraperitoneally (*ip*) or *id* to the contralateral inLN (contra *id*). Compared to systemic administration (*ip*), contra *id* resulted in a survival benefit and a higher number of complete responders (CR) in CLND mice (**Fig. 6F**). These results suggest that delivering ICB therapeutics to non-tdLNs may benefit patients after LN surgery.

In comparison to adjuvant-only treatment, neoadjuvant-adjuvant ICB has been shown to be beneficial for patients with advanced melanoma^50^. To examine whether LN surgery compromises neoadjuvant-adjuvant ICB efficacy, B16F10^Gluc^ tumors were implanted, and mice were treated with ICB delivered *id* to the tdLN followed by a curative primary tumor resection with or without LN surgery (SLNB or CLND). Mice were monitored for tumor recurrence or metastases (using Gluc activity measurement) for 100 days. All tumor-free complete responders were rechallenged with B16F10^Gluc^ tumors and mice that developed tumors were again treated with adjuvant ICB delivered *id* to the ipsi inLN for sham, ipsi axLN for SLNB or contra inLN for CLND (**Fig. 6G**). Interestingly, vitiligo occurred in mice after tumor/LN surgery with neo-adjuvant ICB treatment, which was primarily located at the site of surgery of the tumor resection and LN removal (**Fig. 6H**). Vitiligo indicates an ICB-induced immune response against proteins involved in melanin production beyond the melanin expressing tumor cells^51^. After neoadjuvant treatment and tumor resection, one sham mouse developed a spontaneous tumor recurrence and one mouse of the SLNB group died due to tumor-unrelated reasons (**Fig. 6I**), demonstrating the potency of curative surgery. Upon rechallenge, all naïve control mice developed tumors, while many of the ICB-treated mice rejected the new tumor, indicating anti-tumor immune memory that was more abundant in mice with ICB treatment compared to the IgG controls. The ICB-induced immune memory and survival benefit persisted after SLNB or CLND (**Fig. 6I,J**). These results suggest that, in a LN-targeted neoadjuvant-adjuvant ICB setting, the formation of immune memory against cancer cells and improvement of overall survival persist after LN surgery and curative primary tumor resection.

### LN egress of T cells is dispensable for effective anti-tumor immune memory

ICB generated complete responders in mice with sham surgery or CLND (**Fig. 2**, **Fig. 6**). These complete responders were rechallenged with E0771 tumors at the initial tumor site and were rejected in both groups (**Figure S7B**). We further tested whether the anti-tumor immune memory is contained only in the local tissue or is systemic. All animals rechallenged in both the ipsi- and contra-lateral sides (*bilateral*) remained tumor-free. We further found that the immune memory is tumor specific, as E0771-immune mice did not reject B16F10 tumors implanted in either the mammary fat pad or dermis (**Figure S7B**). Removing the spleen did not influence tumor rejection in complete responders (**Figure S7C**), whereas depletion of CD8 T cells during tumor rechallenge abolished anti-tumor immune memory (**Figure S7D**). Lymphocytes, including CD8 T cells, use sphingosine 1-phosphate (S1PR) as guidance cues to egress from LNs^52–54^. FTY720, a potent inhibitor of S1PR^55^, drastically reduced CD4, CD8 and central memory (CD44^+^/CD62L^+^) CD8 T cells within the circulation (**Figure S7E**). Inhibiting T cell egress from LNs with FTY720 abrogated ICB responses completely during initial tumor growth (**Figure S7F**) prompting us to test whether central memory T cells from LNs enter the circulation as contributors to the long-term immune memory in complete responders. Tumor naive mice (**Figure S7G)** or tumor-free survivors (**Figure S7H, I)** were treated with FTY720, and E0771 or B16F10^Gluc^ tumors were orthotopically implanted to the primary tumor site and to the contralateral side. FTY720 inhibited ICB responses in the tumor naive mice (**Figure S7G).** The vast majority of ICB complete responders rejected the tumors after rechallenge regardless of the FTY720 treatment or extent of initial LN surgery (**Figure S7H, I**). Strikingly, the anti-tumor immune memory was not only effective in rejecting the tumor at the initial tumor implantation site, but also for tumors implanted ectopically (dermis for E0771 and mammary fat pad for B16F10^Gluc^) (**Figure S7H,I)**. These findings suggest that ICB-induced anti-tumor immune memory provides systemic immunity against tumor rechallenge in complete responders while immune cells egressing from LNs are dispensable once the tumor-specific immune memory has been established. Resecting the primary tdLNs, via SLNB or CLND after ICB treatment, had no detrimental effects on tumor rejection mediated by memory T cells.

### Distant LNs become reactive after tumor and tdLN resection in patients treated by ICB

To examine whether non-tdLNs contribute to ICB efficacy against cancer recurrence, we investigated patients from a recent phase I/Ib trial (NCT03635164)^56^. The trial included patients with HPV-negative head and neck cancer who received neoadjuvant treatment consisting of a combination of hypofractionated radiotherapy and durvalumab (targeting PD-L1). After a recovery time, patients underwent surgical resection of the primary tumor and bilateral LN dissection in the neck as standard of care. Patients with a major or complete pathological response to neoadjuvant therapy were classified as responders (response>95%). Except for four cycles of durvalumab treatment, other adjuvant therapies (e.g. radiation- or chemo-therapy) were omitted in responders^56^. No recurrent disease was detected within the responder group during post-operative follow-up visits (>2 years). We compared CT and PET scan images at the time of the diagnosis and post-surgery follow-up examinations. Strikingly, after removal of the tumor and tdLNs, distant LNs with clear signs of expansion/reactivity were found in most responders, while non-responders lacked reactive distant LNs to the initial site of the tumor (**Fig. 7A,B**). Of note, no metastatic LN involvement was detected in radiology-guided biopsies, indicating that enlarged LNs were indeed immune-reactive. The reactive LNs were higher in the echelon and likely located in new lymph draining routes of the initial tumor site after the removal of the primary tdLNs. For instance, in two patients (#13, #15), lower neck LNs (posterior cervical) and upper neck LNs (submandibular and submental) were enlarged during the follow-up though unremarkable at the time of diagnosis (**Fig. 7C**). Similarly, neck draining mediastinal LNs above the aortic arch in patient #9 were normal at the time of diagnoses but appeared reactive in CT and PET scans 7-month after tumor and tdLN dissection (**Fig. 7D**). These data suggest that surgical removal of the tdLNs results in a compensatory mechanism that includes antigen diversion to distant LNs that extends ICB response in cancer patients.

**Figure 7.**
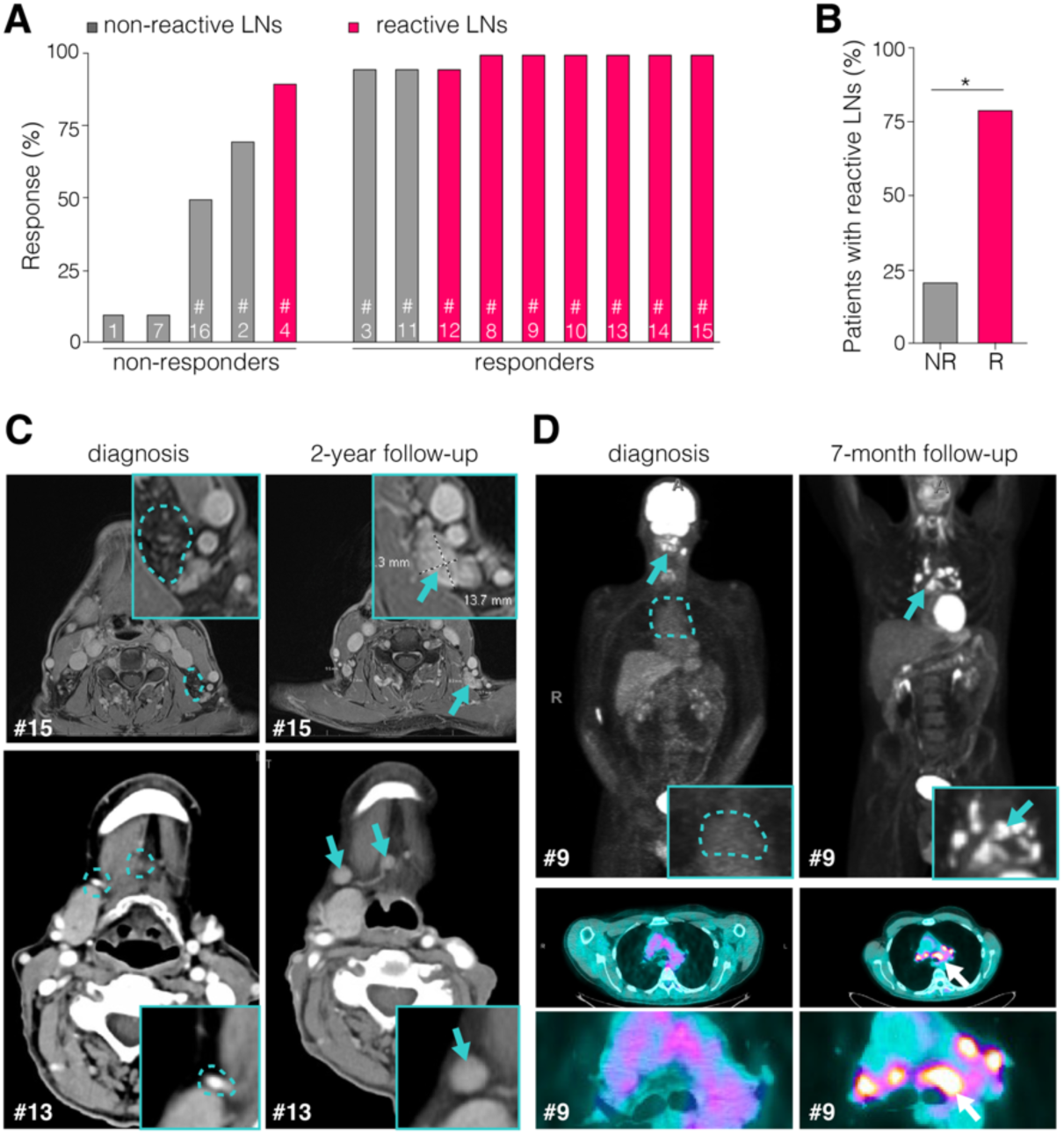
Non-tdLNs become reactive after resection of the primary tumor and related LNs in cancer patients. **(A)** Response rate and reactive status of remaining LNs after primary tumor and LN dissection in the neck of patients with head and neck cancer. **(B)** Percentage of patients with enlarged or reactive LNs in more distant LN basins. **(C)** CT scan images of patients #13 and #15 at the time of diagnosis and at follow-up 2-years post-surgery. Arrows indicate enlarged neck LNs (posterior cervical LNs in patient #13; submandibular and submental LNs in patient #15). **(D)** CT and PET scan images of patient #9 at diagnosis and at follow-up 7-month post-surgery. Arrows indicate reactive cervical (diagnosis) and apex mediastinal LNs (follow up). All data and images are part of the phase I trial NCT03635164. Statistical analysis performed with Fisher’s exact test (B), \**p*<0.05.

## Discussion

TdLNs are located along the lymphatic drainage pathway of primary tumors and have the highest concentration of tumor-derived antigens, which renders them the primary sites to initiate and sustain tumor-specific T cell responses^17,57,58^. Patients with melanoma from a phase II clinical trial with neoadjuvant-adjuvant anti-PD1 (pembrolizumab) treatment had improved event-free survival compared to the adjuvant-only setting^50^, indicating that using the primary tumor as the source of antigen and the tdLNs for tumor-specific T cell priming before primary tumor resection is more effective in establishing ICB responses. Recent studies advocate that tdLNs are indispensable for acute and enduring T memory-driven ICB responses^25,26,32^. However, there are data demonstrating that patients still benefit from ICB treatment after sentinel LN and bulky disease-related LN removal^33,59–61^, which is in line with our retrospective analysis of stage III melanoma patients (**Fig. 1**). Collectively, these data demonstrate that patients benefited from adjuvant ICB with no significant effect of tdLN removal surgery. Recent work from our lab and others has shown that metastatic tdLNs are immune suppressed and can impair systemic anti-cancer immune responses^14–17^. Ultimately, the metastatic burden in tdLNs may determine whether SLNB or CLND has a positive, negative or neutral effect on disease recurrence and survival. As human LN chains are complex with multiple echelons^62^, they determine early metastatic involvement and immune activation or suppression status of the sentinel LNs and LNs further downstream. When dissection of metastatic LNs—which are likely to be severely immunosuppressed^63^ and restricted in their ability to mount effective T cell responses^64^—is performed, the net benefit of removing these immune suppressive LNs is likely to outweigh any marginal decrease in systemic response to ICB. Based on the local microenvironment experienced by individual LNs— including the presence of metastatic cancer cells, abundance of cancer antigen and immunosuppressive cytokines—different tdLNs can be expected to have different capacities to mount immune responses and maintain responses to ICB. It is key to identify LNs with sufficient exposure to antigen and efficient antigen presentation without immune suppression to target for immunotherapy delivery.

In this study, we emphasize the dynamic and adaptive aspects of antigen transport. Disruption of normal lymphatic drainage diverts interstitial fluid with antigens to adjacent lymphosomes and induces immune responses in non-tdLNs. These LNs can be relatively distant from the original tdLNs, which seems to be counterintuitive but is corroborated by clinical observations^42,65^ (**Fig. 7)**. It is likely that the amount/concentration of tumor-derived antigen will still be critical to meet a threshold for effective immune activation after the transport distance to distant LNs. Long distance transport goes along with antigen dilution and degradation and might fail to induce the inflammatory milieu in LNs that is required for proper T cell priming. Understanding these spatial limits might explain the anatomical distribution of LNs throughout the body needed to provide effective regional immune surveillance. Intriguingly, we found that CLND in mice induces tumor-specific T cell enrichment in distant LNs with ICB treatment, when performed along with primary tumor resection (adjuvant setting). The antigen source fueling the tumor-specific T cell response in distant LNs after primary tumor resection is unknown. Conceivably, surgery might release a larger amount of antigen. Residual tumor cells from the primary site or metastases in distant organs can also contribute to the antigen abundance. Despite the importance of antigen transport to distant LNs, direct T cell priming in the local tumor microenvironment^66^ and tertiary lymphoid structures^67^ may also contribute to the residual ICB response.

Our findings suggest that long-term anti-tumor immune memory is not predominantly reliant on T cell egress from LNs. It is possible that tumor-specific memory T cells leave the tdLNs after initial activation and reside in tissues until getting reactivated during tumor regrowth or metastasis. Our neoadjuvant-adjuvant ICB study demonstrated that mice that underwent curative surgery retained tumor-specific systemic immune memory, suggesting that adjuvant ICB may act through existing anti-tumor tissue-resident memory T cells.

In our work, surgery of skin LNs in mice was used to recapitulate SLNB and CLND procedures used in the clinic. However, the length scale and anatomical differences are obvious between humans and mice (**Fig. 4)**. Compared to simple chains of a few LNs in a drainage path of mice, humans have more complex and interconnected lymphatic networks with multiple LNs draining the same lymphosome^62^, so that the removal of a single LN in mice resembles CLND in humans. Consistent with our preclinical data, compensatory lymphatic drainage^42,65^, even in the presence of lymphedema^40–42,65^, and distant LN activation are observed in cancer patients (**Fig. 7)**. In future work, it will be of interest to investigate lymphatic drainage patterns apart from the lymphatic network of the skin, as ICB treatment also increases survival in non-small cell lung cancer^68^ and colorectal cancer^69^, which may have less access to adjacent lymphosomes to access alternative LNs. Our data present a better understanding of the challenges associated with integrating ICB therapy into clinical practice in patients undergoing LN resections. It is critical to continue to advance this understanding to improve ICB responses in patients with metastatic cancer.

## Supplemental Figures

**Figure S1.**
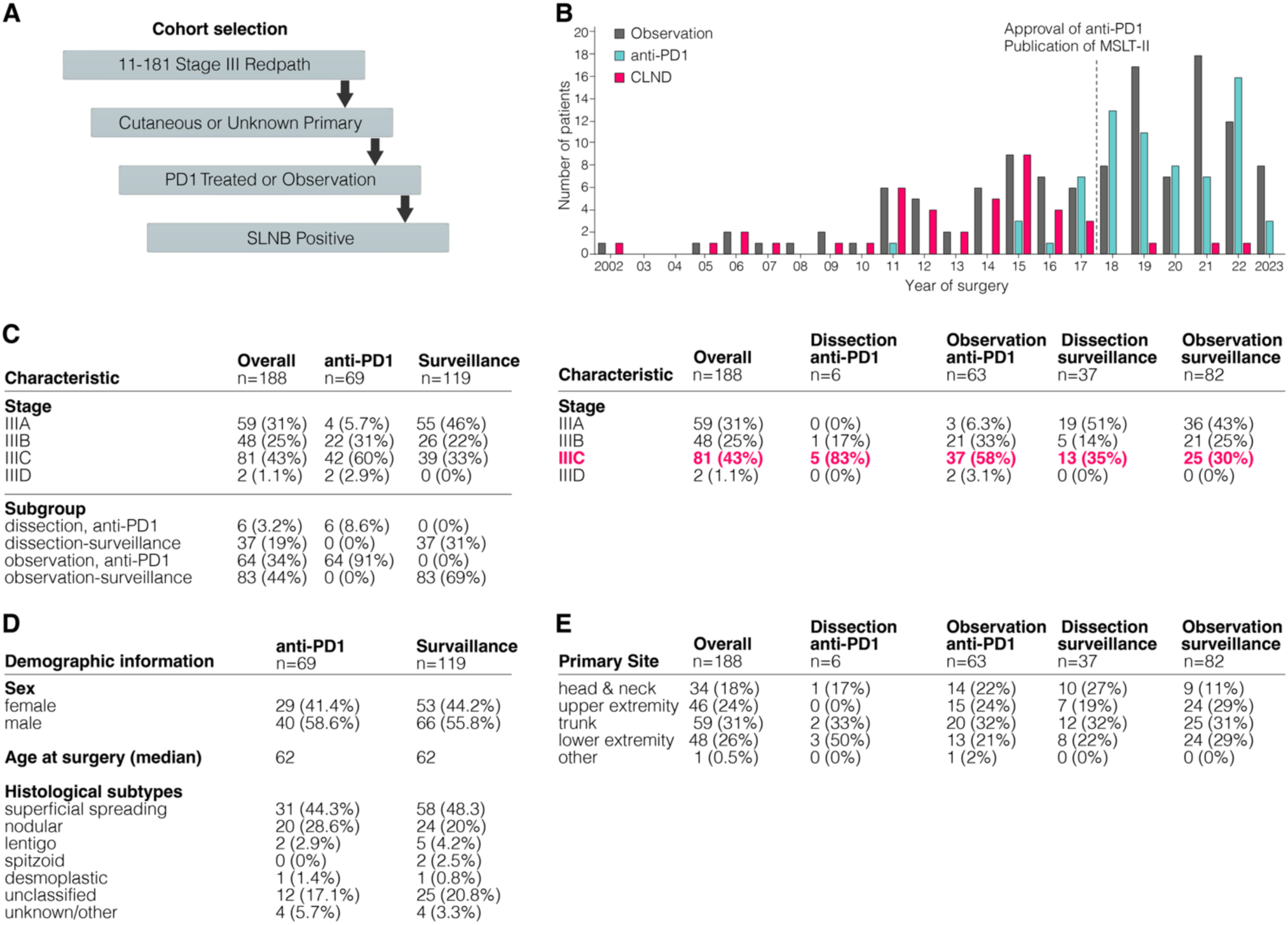
**(A)** Cohort selection strategy. **(B)** Historical changes of the number of patients untreated (observation), treated with anti-PD1 therapy or LN surgery (CLND) in combination with adjuvant therapy regimen. MSLT-II and anti-PD1 trials indicated on timeline as major milestones. **(C)** Composition and subgrouping of patients. **(D)** Demographic information and **(E)** information about the primary tumor site of patients with LN surgery and surveillance or anti-PD1 treatment.

**Figure S2.**
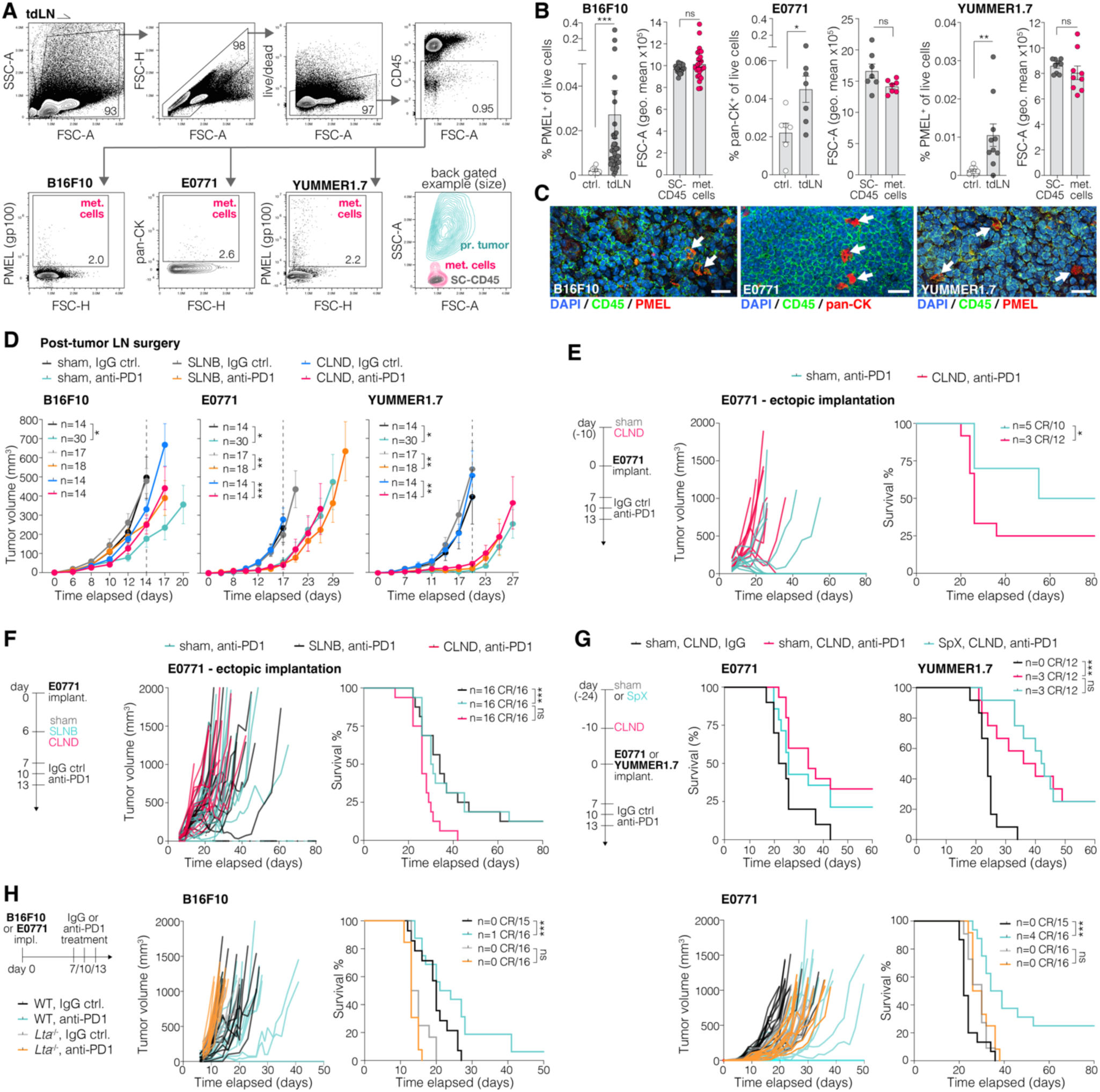
**(A)** To evaluate the occurrence of LN metastases with the experimental set up used in the study, B16F10, E0771 or YUMMER1.7 tumors were orthotopically implanted and LNs analyzed for tumor cells by flow cytometry. **(B)** Frequency of tumor cells relative to live cells and size of events (geometric mean of FSC-A). **(C)** Representative pictures of tdLN showing immune cells (CD45^+^) and metastatic tumor cells (PMEL^+^ or pan-cytokeratin^+^ (pan-CK). Scale bar: 15 µm. **(D)** Average tumor growth curves of post-tumor LN surgery, corresponding to Fig. 2A. Dotted grey line indicates time point of statistical analysis between groups. **(E,F)** Ectopically implanted (intradermal in the flank) E0771 tumor growth and survival with (E) preemptive sham surgery or CLND or (F) post-tumor sham surgery, SLNB or CLDN. **(G)** Growth and survival of E0771 or YUMMER1.7 tumors implanted in mice after sham surgery or splenectomy (SpX) in combination with CLND and IgG control or anti-PD1 treatment. **(H)** Growth and survival of B16F10 or E0771 tumors implanted in WT or *Lta*^−/−^ transgenic mice and treated with IgG or anti-PD1 antibody. WT mice survival data are from sham IgG and sham anti-PD1 conditions from Figure 2B for comparison. Number of complete responders (CR) of total mice are indicated. Graphs show mean ± SEM with data points representing individual animals (B). Statistical analysis with Mann-Whitney U test (B), 2-way ANOVA (D) or Log-rank test (E,F,G,H), ns= not significant, \**p*<0.05, \*\**p*<0.01, \*\*\**p*<0.005.

**Figure S3.**
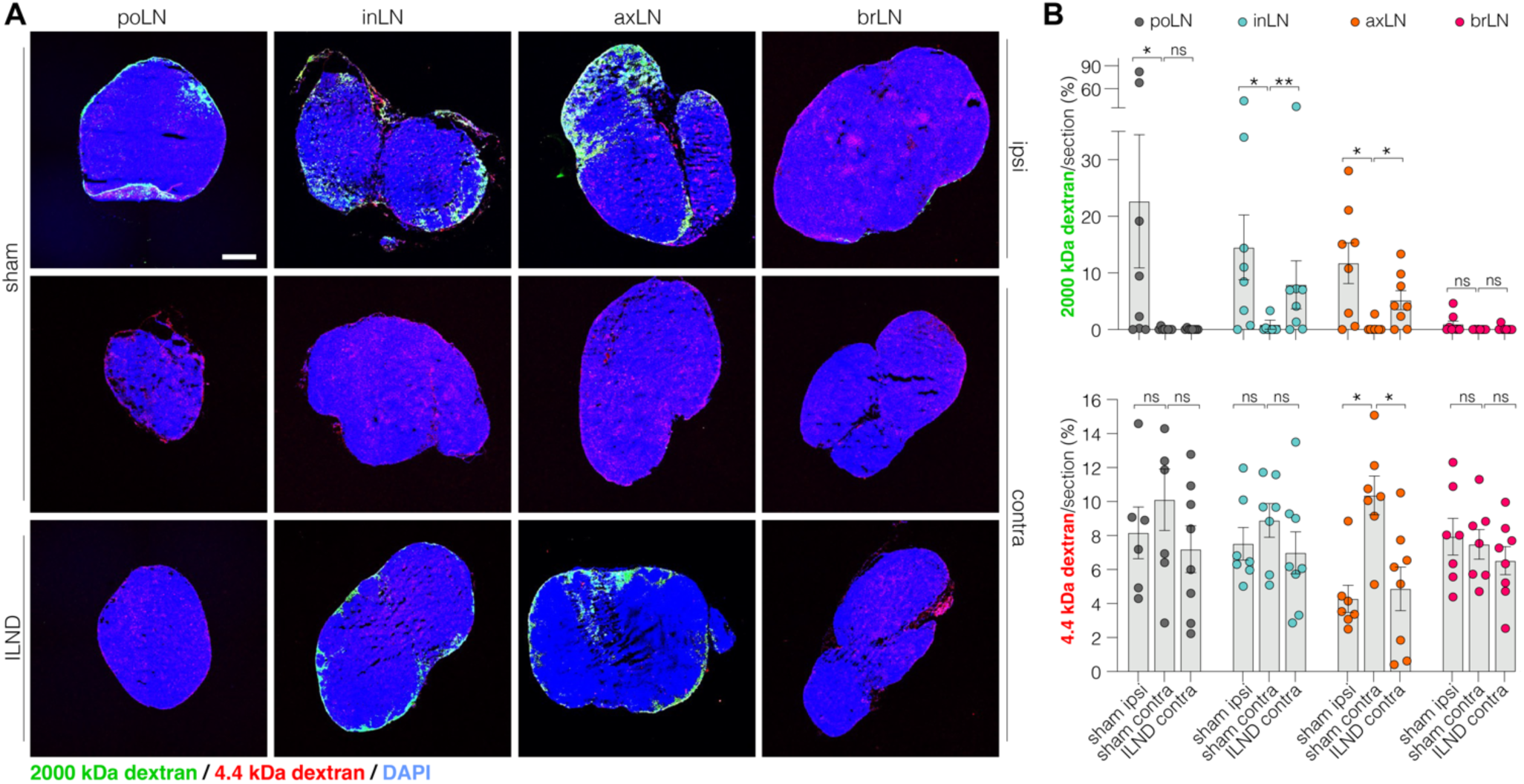
LN dissection promotes antigen drainage to distant LNs. FITC-conjugated 2000 kDa dextran and TRITC-conjugated 4.4 kDa dextran were injected to the footpad of mice after sham surgery or ipsilateral LN dissection (ILND, including the popliteal (poLN), inguinal (inLN), axillary (axLN) and brachial (brLN)). **(A)** Representative images of dextran distribution in the poLN, axLN, inLN and brLN. Scale bar: 200 µm. **(B)** Quantification of the fluorescent area (FITC or TRITC) in percentage of whole section. Graphs show mean ± SEM with data points representing individual animals. Statistical analysis with MWU test, ns= not significant, \**p*<0.05, \*\**p*<0.01.

**Figure S4.**
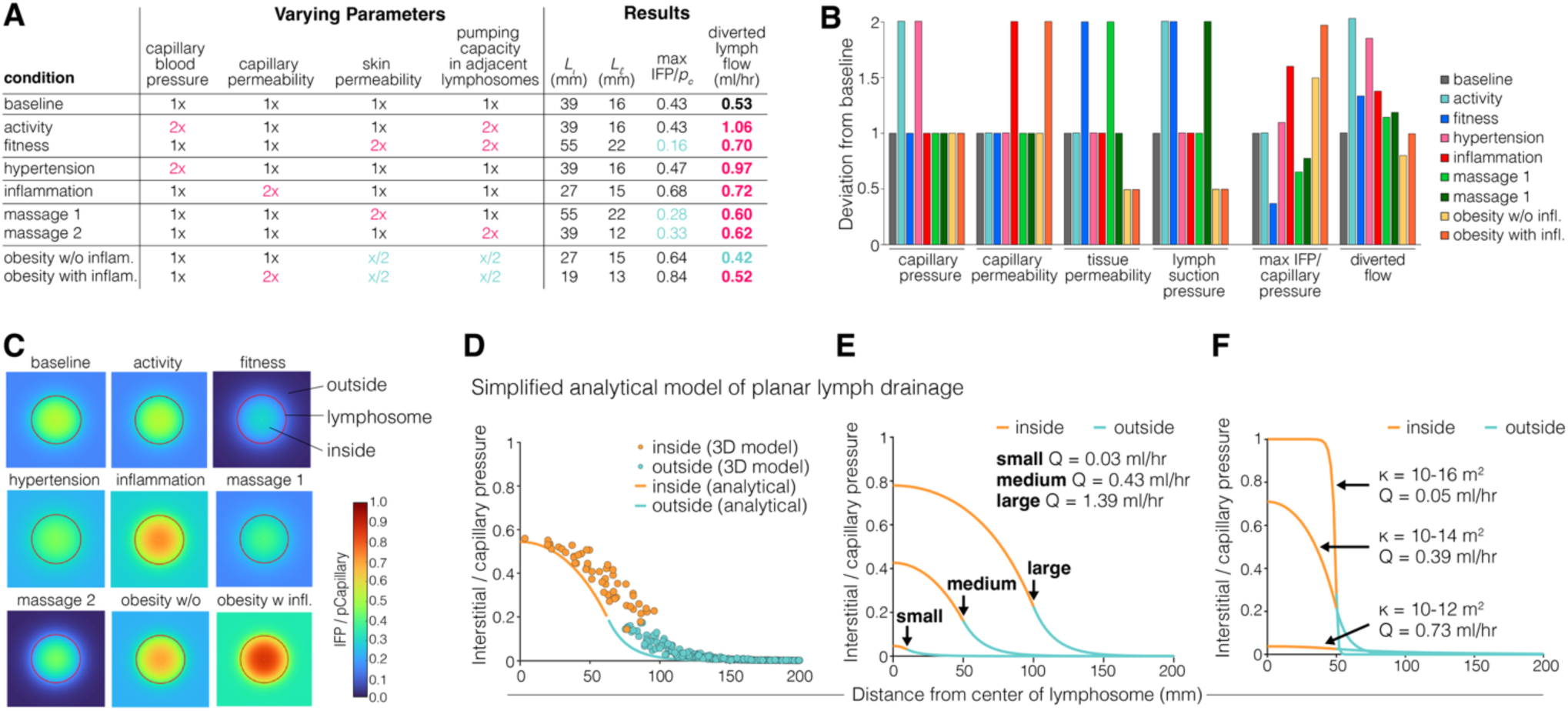
Varied parameters during different body conditions with simplified analytical model of planar lymph drainage. **(A)** Varied parameters during different body conditions. Physical *activity* is represented by a twofold increase in both capillary blood pressure and lymphatic suction, *fitness* achieved through extended periods of activity by twofold increases in tissue permeability and lymph suction pressure, *hypertension* by increased capillary pressure, widespread *inflammation* by increased capillary permeability, *massage* by increased skin permeability or suction pressure, and *obesity* by halved tissue permeability and lymph suction pressure which may be combined with inflammation. The results show that fitness and massage tend to offset the effects of obesity by reducing interstitial pressure while increasing the quantity of diverted fluid. **(B)** Deviation from baseline corresponding to the conditions in (A). **(C)** Interstitial fluid pressure relative to the capillary blood pressure mapped for four, approximate physiological conditions for a 50 mm radius lymphosome (red circle). **(D)** Dependence of interstitial pressure inside and outside the disabled lymphosome from the full human model in Figure 4I compared to the analytical prediction for a circle of the same area. **(E)** Dependence of interstitial pressure on the radius of the disabled lymphosome (arrows) and rate of fluid diversion (Q) at the periphery of the disabled lymphosome showing that small lymphosomes produce little increase in interstitial pressure and that the quantity of diverted flow grows in rough proportion to the area of the disabled lymphosome. **(F)** Dependence of interstitial pressure and rate of fluid diversion (Q) at the periphery of the disabled lymphosome (radius 5 mm), showing that changes in tissue permeability alter the interstitial pressure, but only weakly alter the rate of fluid/antigen flow towards compensatory lymphosomes. Baseline parameters in supplemental methods.

**Figure S5.**
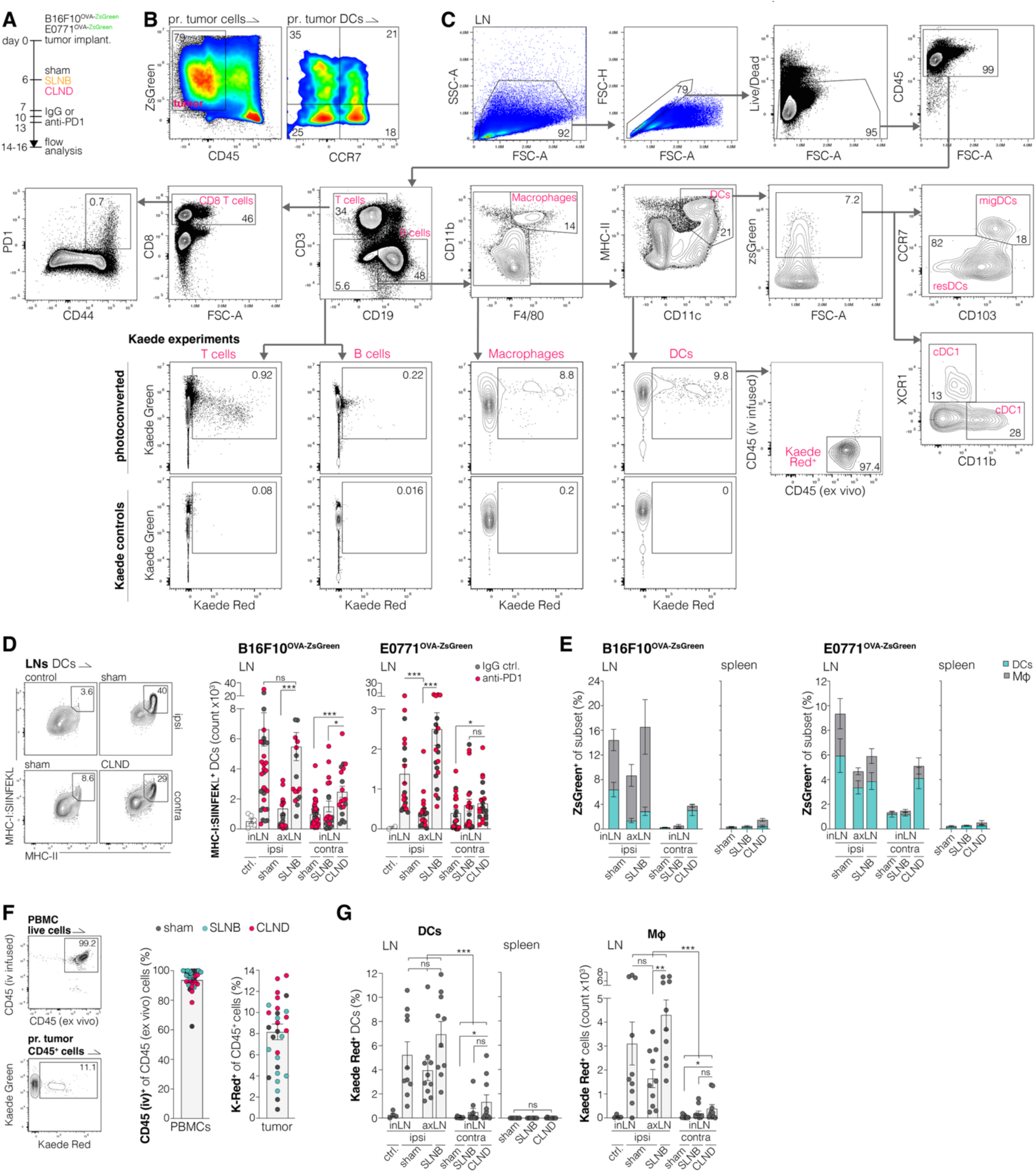
**(A)** Experimental set up to analyze DC migration and tumor-derived antigen presentation in LNs. Corresponding to C, D, E; Fig. 5A,B,C and Fig. 6A. **(B)** Representative flow cytometry plots of ZsGreen^+^ tumor cells and DCs within the primary tumor. **(C)** Gating strategy for flow cytometry analysis of LNs and spleen, including Kaede experiments with controls (*lower panel*). **(D)** Flow analysis of DCs presenting SIINFEKL via MHC-I (antibody clone 25-D1.16), including IgG control and anti-PD1 treated animals. Controls represent mice without tumor implantation. **(E)** Percentage of ZsGreen^+^ DCs and macrophages (MΦ) in LNs and spleen. **(F)** Representative plots and quantification of PBMCs from Kaede mice intravenously (iv) infused and *ex vivo* stained with CD45 antibodies (upper plot). Photoconverted CD45^+^ cells within the primary tumor at the time of analysis (lower plot). **(G)** Percentage of Kaede^+^ DCs and macrophages (MΦ) in LNs and spleen show limited migration to the contralateral LNs. Graphs display mean ± SEM, data points represent individual animals of at least two independent experiments. Statistical analysis with Mann-Whitney U test (D,G), ns= not significant, \**p*<0.05, \*\**p*<0.01, \*\*\**p*<0.005.

**Figure S6.**
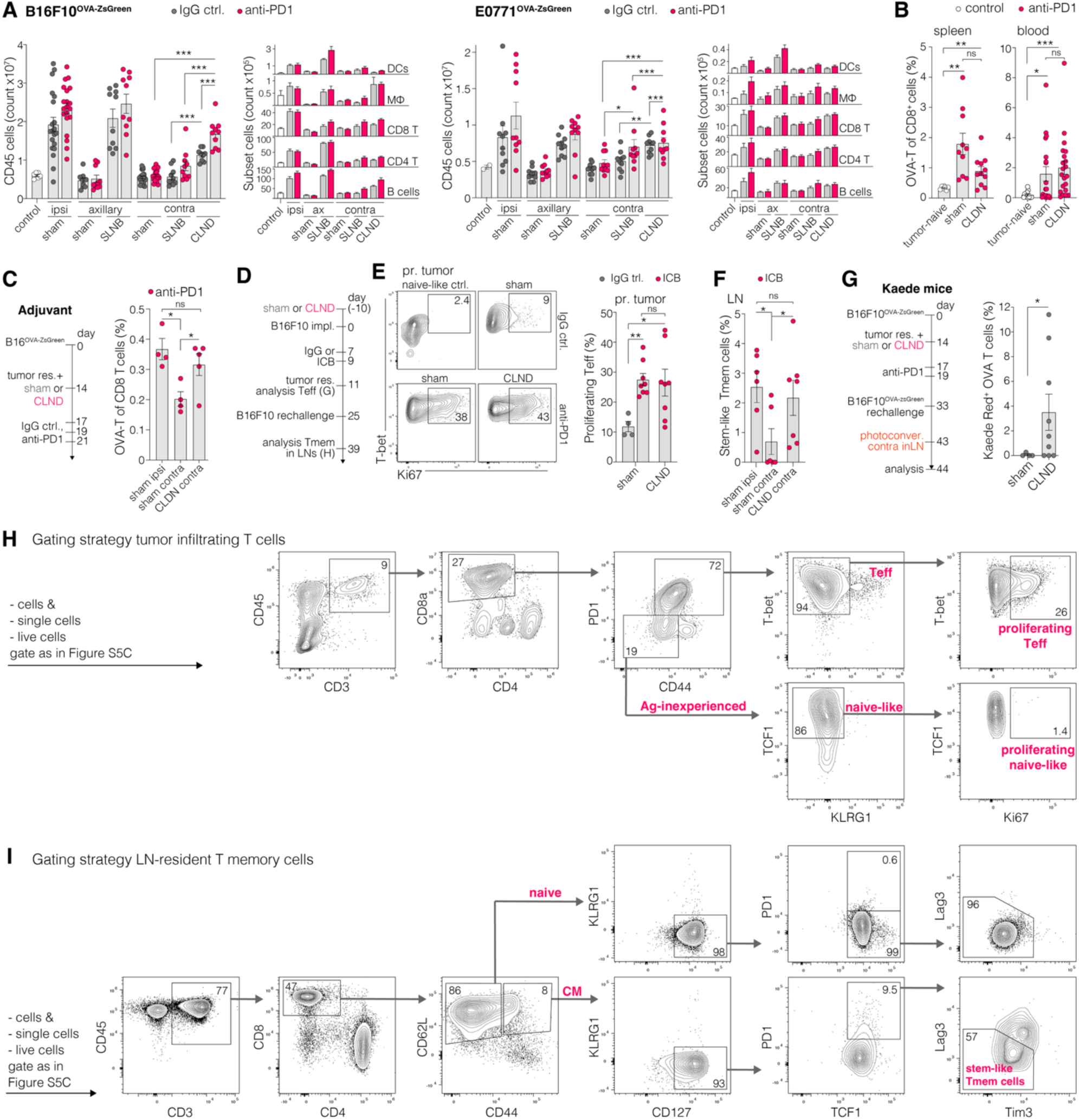
**(A)** LN cellularity of CD45+ leukocytes and immune composition. Corresponding to Fig. 5A-C and Fig. 6A. Control represents mice without tumor implantation **(B)** OVA CD8 T cells in spleen and blood of mice with B16F10^OVA-ZsGreen^ tumors. Corresponding to Fig. 6B. **(C)** Frequency of OVA-specific CD8 T cells in LNs in an adjuvant setting with primary tumor resection combined with CLND and anti-PD1 treatment. **(D)** Experimental set up to analyze effector T cells (Teff) in the primary tumor and memory T cells (Tmem) in LNs after tumor rechallenge. Corresponding to E, F. **(E)** Proliferating CD8 effector T cells within the primary tumor. **(F)** Stem-like T memory cells in LNs of mice after tumor rechallenge. **(G)** Experimental set up to analyze trafficking of OVA-T cells from photo-converted contralateral LNs into the primary tumor after tumor rechallenge in Kaede mice. Control animals had the primary tumor resected without LN dissection. Graph shows the number of KaedeRed OVA-T cells from the photoconverted contralateral inguinal LN that were found in the rechallenge tumor. **(H)** Gating strategy to analyze tumor infiltrating T cells and **(I)** LN-resident T memory cells. Statistical analysis with Mann-Whitney U test (A,B,C,E,F), Welch’s test (G), \**p*<0.05.

**Figure S7.**
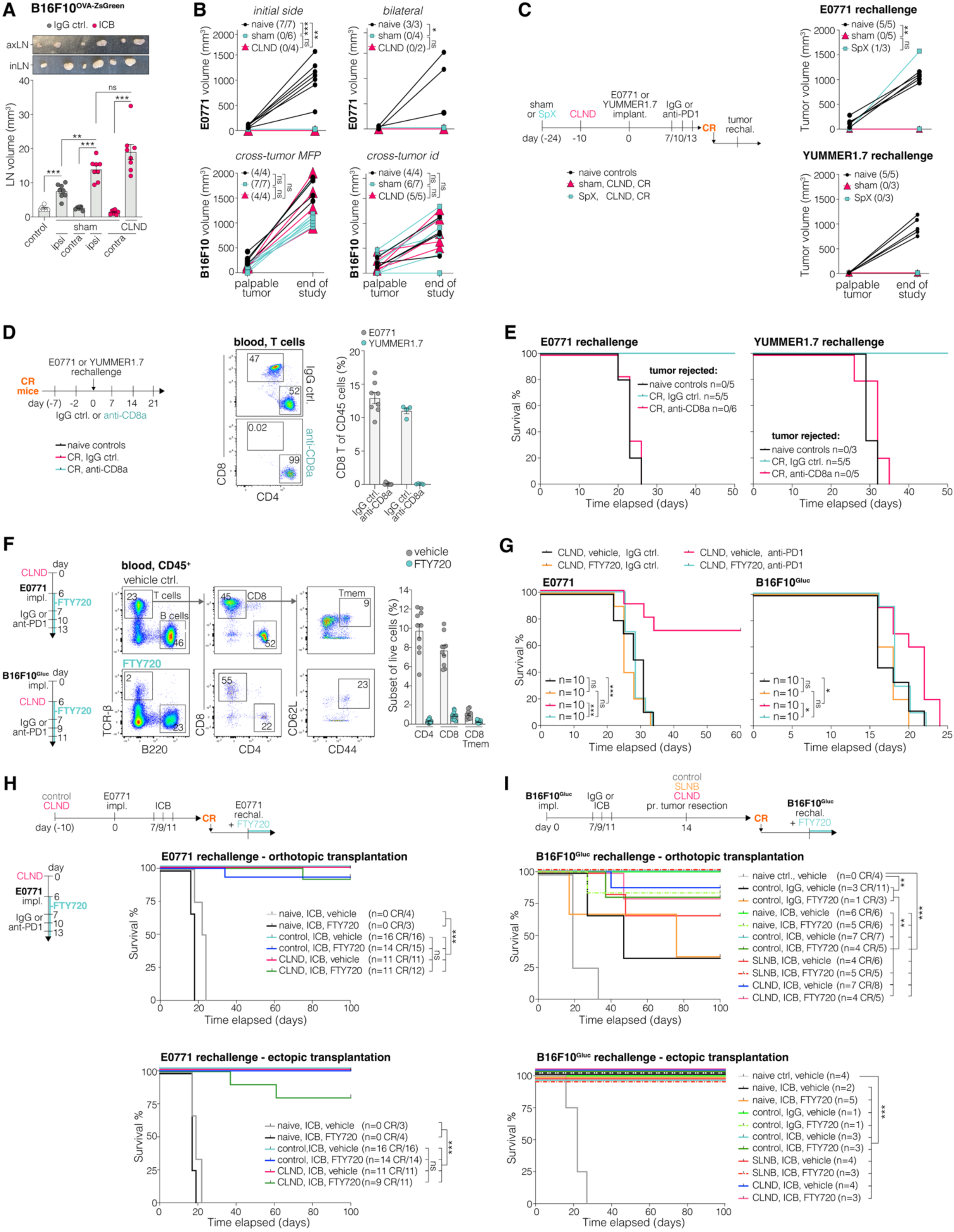
Acute T cell egress from LNs is dispensable for long-term anti-tumor immune memory. **(A)** LN volume in mice with B16F10^OVA-ZsGreen^ tumor that underwent sham or CLND surgery followed by IgG or the combination of anti-CTLA4 and anti-PD1 (ICB) treatment. Corresponding to Fig. 6C**-E**. **(B)** Growth of E0771 tumors in naive mice or in complete ICB responders implanted at the primary tumor site (*upper left*), or both ipsilateral (primary tumor site) and contralateral side (*upper right*). B16F10 tumors implanted in the mammary fat pad (*lower left*) or dermis (*lower right*) of naive mice or complete responders after E0771 tumor rejection. The number of animals with palpable tumors in parenthesis in the figure legend. **(C)** E0771 or YUMMER1.7 tumors implanted in complete responder (CR) mice after sham surgery or splenectomy (SpX) in combination with CLND and IgG control or anti-PD1 treatment. Corresponding to **Figure S2E**. **(D)** *Left*, Experimental set up and, *right*, Representative flow cytometry plots and quantification demonstrating the depletion of CD8 T cells in blood samples of mice treated as indicated with IgG control or anti-CD8a antibodies. **(E)** Survival of animals with E0771 or YUMMER1.7 tumor rechallenge in complete responders (CR) with CD8 T cell depletion or IgG controls. Tumor naive animals were used as untreated controls. **(F)** *Left*, Experimental set up and, *right*, Representative flow cytometry plots and quantification demonstrating the abundance of circulating B cells, T cells and central memory T cells in the blood after FTY720 added to the drinking water. **(G)** Survival of animals with E0771 or B16F10^Gluc^ tumor implantation and IgG control or anti-PD1 treatment while the lymphocyte egress from LNs is inhibited with FTY720. **(H)** Survival of previous long-term survivor animals that underwent pre-emptive LN surgery and ICB treatment that were subsequently rechallenged with E0771 in an orthotopic site (MFP) or ectopic site (dermis) sequentially on both sides and treated with vehicle or FTY720. Tumor-naive mice were used as naive controls, control group represents tumor resection without LN surgery. **(I)** Survival of previous long-term survivor animals that underwent neo-adjuvant ICB and B16F10^Gluc^ tumor rechallenge to orthotopic site (dermis) or ectopic site (MFP) sequentially on both sides and treated with vehicle or FTY720. Graphs show mean ± SEM, data points representing individual animals. Statistical analysis with Mann-Whitney U test (A), Fisher’s exact test (B,C) or Log-rank test (E,G,H,I), ns= not significant, \**p*<0.05, \*\**p*<0.01, \*\*\**p*<0.005.

## Resource Availability

Lead contact: Requests for further information and resources should be directed to and will be fulfilled by the lead contact, Timothy P. Padera

Material availability: This study did not generate new unique reagents or mouse lines.

Data and code availability: All data reported in this paper will be shared by the lead contact upon request.

## ACKNOWLEDGEMENTS

We thank Peigen Huang and the Cox-7 animal facility for maintaining and providing mice and technical support, Mark Duquette for providing help and technical assistance, and Stefanie Spranger for providing plasmids. We thank the NIH Tetramer Core Facility (contract number 75N93020D00005) for providing tetramers for flow cytometry.

## Funding

The Rullo Family MGH Research Scholar Award (T.P.P.)

Emma and Bill Roberts MGH Research Scholar Award (G.M.B.)

National Institutes of Health grant K00CA234940 (H.Z.)

National Institutes of Health grant R21AG072205 (T.P.P.)

National Institutes of Health grant R01CA214913 (T.P.P.)

National Institutes of Health grant R01HL128168 (T.P.P.)

National Institutes of Health grant R01CA284372 (T.P.P.)

National Institutes of Health grant R01CA284603 (T.P.P., L.L.M. and J.W.B.)

National Institutes of Health grant R01DE028529 (S.D.K.)

National Institutes of Health grant R01DE028282 (S.D.K.)

National Institutes of Health grant P50CA261605 (S.D.K.)

National Institutes of Health grant R35CA242379 (M.G.V.H.)

The Bridge Project, a partnership between the Koch Institute for Integrative Cancer Research at MIT and the Dana-Farber/Harvard Cancer Center (T.P.P., M.G.V.H.)

The Patricia K. Donahoe Award from the Huiying Foundation (G.M.B.)

The Adelson Medical Research Foundation (G.M.B.),

The Ludwig Center at MIT (M.G.V.H.)

The MIT Center for Precision Cancer Medicine (M.G.V.H.)

The Breast Cancer Alliance (J.M.U.)

The Melanoma Research Foundation (J.M.U.).

Deutsche Forschungsgemeinschaft ME5486/1-2 (L.M.)

## Author Contributions

Conceptualization: H.Z., L.M., J.W.B., M.G.V.H., L.L.M., G.M.B., S.C., S.D.K., T.P.P.

Methodology: H.Z., L.M., J.W.B., M.G.V.H., L.L.M., G.M.B., S.C., S.D.K., T.P.P.

Data acquisition: H.Z., L.M., J.W.B., M.J.O., L.B.D., E.S., J.C., P.J.L., J.J.R.

Intellectual input: H.Z., L.M., J.W.B., M.J.O., P.J.L., M.R.N., M.S.R., M.G.V.H., J.M.U., L.L.M., G.M.B., S.C., S.D.K., T.P.P.

Project administration: H.Z., L.M., T.P.P. Supervision: G.M.B., S.C., S.D.K., M.G.V.H., T.P.P.

Writing: H.Z., L.M., J.W.B., M.J.O., P.J.L., M.G.V.H., J.M.U., L.L.M., G.M.B., S.C., S.D.K., T.P.P.

