## Supplemental Methods for "Distant lymph nodes compensate for resected tumor-draining lymph nodes during cancer immunotherapy"

### MATERIALS AND METHODS

#### Retrospective analysis of melanoma patient survival

We performed a retrospective analysis of patients with stage III cutaneous melanoma according to the $8^{th}$ American Joint Committee on Cancer criteria. The patients were selected from a large melanoma cohort study and tumor biobank maintained at Massachusetts General Hospital for over 13 years (2009-2022) and approved by the Institutional Review Board. Informed consent was obtained from all patients. All patients included in this analysis received at least one line of anti-PD1 immune checkpoint blockade adjuvant therapy. Among patients with SLNB+ diagnosis, only the stage IIIC disease subcohort was selected for analysis, to fairly compare all surgery and treatment conditions. For each patient, we extracted general demographic information, histopathological reports, extent of surgery, adjuvant treatments, recurrence and survival outcomes. The data extraction was followed by statistical analysis using R (Core Team 2021) package(75) (survminer) to generate Kaplan-Meier survival curves. Significance was determined using a log-rank test and the p-value is reported on each plot. P-value significance was considered to be $p<0.05$.

#### Clinical trial data and patient records

In phase I/Ib trial NCT03635164 pathological response was evaluated independently by two board-certified pathologists who were blinded to treatment outcome. Specimens collected at time of surgery were evaluated for microscopic residual tumor (only scattered residual foci) or macroscopic residual tumor and scored based on these findings.

In phase I/Ib trial NCT03635164, patients received a baseline PET/CT of skull base to mid-thigh and an MRI with and without contrast of the neck no more than 28 days before radiation therapy. Long-term follow up imaging assessment was done with either PET/CT and/or MRI at the discretion of the treating physician. Enlarged lymph nodes at the follow-up timepoints were noted on imaging by a radiologist and concerning nodes were biopsied to rule out recurrence or otherwise considered to be reactive.

Reactivity was defined by board certified neuroradiologists and nuclear medicine physicians. Biopsies were performed to confirm absence of disease. The cases were discussed at multi-disciplinary tumor board and these patients were followed over time (2.5 years now). Lymph nodes that did not change in shape, size, or level of enhancement by PET or MRI over time were called reactive.

#### Cell lines

Parental E0771 (ATCC, CRL-3461) mammary carcinoma line, and parental B16F10 (ATCC, CRL-6475), YUMMER1.7D5 (Sigma, Cat: SCC244) melanoma lines were cultured using Dulbecco's modified Eagle's medium (DMEM, Corning, Cat#10-014-CV) or DMEM:F12 (for YUMMER1.7, incl. 1% non-essential amino acids) supplemented with 10% (v/v) fetal bovine serum at 5 % $CO_2$ and 37 °C. All cells were authenticated before use and checked for mycoplasma monthly using PCR Mycoplasma Test Kit I/C.

#### Transduction

HEK293 cells were transfected with 1μg of plasmids of gene of interest as well as psPAX2 and VSVG, using 5% Fugene transfection reagent in Opt-MEM (Gibco, Cat#11058-021). Lentiviral supernatant combined with polybrene (8μg/ml) was applied to target cells. pLV-Ef1a-ZSgreen-SIINFEKL-IRES-puro plasmid (gift from Stefanie Spranger lab at MIT) was used to generate ZsGreen and ovalbumin (257-

264) overexpressing cells. CSCW-mbGluc-biotin-IG plasmid (generated by MGH Vector Core) was used to generate Gaussia luciferase expressing cells.

**Animal models**

In addition to wild-type C57Bl6 mice, lymphotoxin alpha knockout mice B6.129s2-Lta$^{tm1Dch}$/J (*Lta*$^{-/-}$), from the Jackson Laboratory (Strain# 002258) and Kaede mice (B6 Tg(CAG-tdKaede)15Utr) were used. All mice were bred and maintained in a pathogen-free environment at facilities of the Massachusetts General Hospital, with 12-hour light/dark cycles at 22°C and 55% humidity in compliance with the institutional regulations. Male and female mice between the age of 6-10 weeks were used in all experiments. Experimental groups were sex-matched or used according to the origin of the tumor cell line. All animal experiment protocols were approved by the Institutional Animal Care and Use Committee (IACUC) of the Massachusetts General Hospital, Boston, MA.

**Surgical procedures**

Primary tumor resection: Under anesthesia, fur at the tumor site was removed and the skin sterilized. The tumor was separated from the underlying fascia and overlying skin by blunt dissection, the tumor resected, and the skin closed with wound clips.

LN dissection: Under anesthesia, fur at the tumor site was removed and the skin sterilized. An incision was made to the skin over the inguinal or axillary region and either just the inguinal LN (SLNB) or the inguinal, axillary and brachial LN (CLND) were exposed by blunt dissection, the LNs resected, and the skin closed with wound clips.

Splenectomy: Under anesthesia, an incision was introduced to the right of the ventral midline (below diaphragm). Splenic arteries and veins were ligated at the splenic hilum with silk suture and the spleen removed. The peritoneum was closed with polydioxanone suture and the skin using wound clips. Mice recovered for at least 14 days prior to subsequent experiments.

**Tumor models**

$2x10^5$ E0771 cells or $5x10^5$ E0771$^{OVA\text{-}zsGreen}$ were injected into the 4$^{th}$ mammary fat pad of female C57BL6 mice, $5x10^5$ B16F10, B16F10$^{OVA\text{-}zsGreen}$, B16F10$^{Gluc}$ and YUMMER1.7 cells were implanted in the dermis (bottom flank) of male or female C57BL6 mice. The tumor volume was monitored at least twice per week and mice were sacrificed when tumor volume exceeded 1000 mm$^3$, any diameter exceeded 15 mm, or ulcerations larger than 4 mm occurred. Tumor rechallenge was performed with mice that rejected the tumor after it was successfully engrafted after implantation (complete responders).

**Photoconversion and processing of Kaede mice**

Tumors were photoconverted using a 405 nm light emitting diode at 11 mW with 10 cm distance to the tissue, for 5 min on 3 consecutive days. During photoconversion, all non-tumor tissue was carefully covered with aluminum foil, precisely fitted to individual tumors to avoid exposure of other regions. 3 min before euthanasia, fluorescence-conjugated anti-CD45 antibodies (3 µg/mouse in PBS) were intravenously (iv) infused in every mouse to distinguish between cells in the blood circulation or LN parenchyma. The iv infusion was checked in blood samples and the photoconversion with CD45 cells from the primary tumor by flow cytometry **(Figure SG)**.

**Antibody and Blocking agent treatment regimens**

Immune checkpoint blockage: Anti-PD1 blocking antibody (10 mg/kg) or anti-PD1 in combination with anti-CTLA4 antibody (5 mg/kg) were administered intraperitoneally (ip) for systemic distribution or intradermally (id) in vicinity of a LN of interest. ICB was usually administered 3 times every 2 to 3 days. For neoadjuvant-adjuvant treatment regimen, the adjuvant ICB was administered when the rechallenge tumors exceeded a size of 100 $mm^3$. Rat and Syrian Hamster IgGs were used as controls.

CD8 T cell depletion: Anti-CD8a blocking antibody was administered ip with a concentration of 10 mg/kg (in 0.9 % NaCl solution) for the first injection followed by 5 mg/kg once weekly. On day 9 post-injection, CD8 T-cell depletion in blood samples was confirmed by flow cytometry **(Figure S7D)**.

FTY720 mediated inhibition of lymphocyte egress from LNs: FTY720 was dissolved and added to the drinking water of mice. Water bottles and compound were exchanged weekly and successful inhibition of lymphocyte egress checked by flow cytometry 7 days after beginning of treatment.

**Dextran tracing**

50 mg/ml TRITC-dextran with 4.4 kDa, mixed with 25 mg/ml FITC-dextran with 2,000 kDa molecular weight in a volume of 20 µL were injected to mammary fat pad, dorsal foot pad or dermis (specified in text). After 6 hours, ipsilateral and contralateral popliteal, inguinal, axillary, brachial LN, and the spleen were harvested and processed for microscopic analysis.

**Bone marrow-derived DC transfer (DC vaccination)**

Bone marrow was flushed out from hind limbs of 7-12 week old C57Bl6 mice and cultured in RPMI media supplemented with 10% FBS, 1% Pen/Strep, 50 µM β-mercaptoethanol, 20 ng/ml GM-CSF and 10 ng/mL IL-4 for 9-14 days to differentiate BMDCs. BMDCs were pulsed with 20 µg/ml $OVA_{257-264}$ for 3 hours together with 2 µg/mL LPS. $5x10^5$ pulsed BMDCs (in 20 µl medium) were injected in the foot pad of mice. After 7 days, LNs were obtained for further analysis.

**Immunofluorescence staining and imaging**

Fresh tissue was fixed in ice-cold paraformaldehyde (4% in PBS, overnight (ON) at 4°C), incubated in 30% sucrose (ON at 4°C) and embedded in OCT compound before freezing and storage at -80 °C. The tissue blocks were cut into 10 µm serial sections using a cryostat. Sections were blocked with 5% normal donkey serum for 1 hour at room temperature (RT) and stained with primary antibodies (Table SM1) in humidity chamber ON at 4°C. The sections were washed and incubated with secondary antibodies for 1 hour at RT. DAPI was added as counterstaining before sections were mounted with Vectashield Antifade Mounting Medium. Sections were imaged using an Olympus IX81 laser scanning confocal microscope with FV10-ASW 4.2 software or a ZEISS Axio scan Z.1 with ZEN 2.3 Slidescan software.

**Tissue and cell preparation for flow cytometry**

Fresh tissue was dissociated and filtered using a 70 µm cell strainer. For analysis of antigen presenting cells, LNs were digested with Collagenase-IV (150 units/mL) and DNase1 (100 µg/mL) for 20 min at 37°C before dissociation. Red blood cells were lysed with ACK lysis buffer for 2 minutes. Samples were stained with viability dye: DAPI or Zombie Violet for 15 mins, blocked by Fc receptor (CD16/32) antibody for 15 mins and stained with primary antibody (or tetramer) for 1 hour at 4°C in the dark, then washed with PBS and fixed (FoxP3 Fix/Perm Kit) or with CytoFix (BD). For intracellular staining, antibodies were incubated in permeabilization buffer for 1 hour at 4°C. Flow samples were analyzed using a Fortessa X20 flow cytometer (BD) or Aurora spectral analyzer (5 laser, Cytek) within two weeks after fixation.

**Table SM1, Antibodies and tetramers.** Flow cytometry (FC), immune fluorescence (IF), concentration used (Conc.), application (Appl.).

| Antigen | Source | Cat# | Conc. | Appl. |
|---|---|---|---|---|
| B220 PE | Biolegend | 103208 | 1:200 | FC |
| B220 purified | Biolegend | 103201 | 1:100 | IF |
| CCR7 PE/Cy7 | Biolegend | 120123 | 1:200 | FC |
| CD11c APC | Biolegend | 117309 | 1:200 | FC |
| CD127 PE/Cy5 | Biolegend | 135015 | 1:200 | FC |
| CD16/CD32 purified | Biolegend | 101301 | 1:100 | IF |
| CD19 Alexa Fluor 700 | Biolegend | 115527 | 1:200 | FC |
| CD3 PE/Dazzle 594 | Biolegend | 100245 | 1:200 | FC |
| CD3 PerCP/Cy5.5 | Biolegend | 100217 | 1:200 | FC |
| CD39 PE/Cy7 | Biolegend | 143805 | 1:200 | FC |
| CD44 Brilliant Violet 750 | Biolegend | 103079 | 1:200 | FC |
| CD44 PE | Biolegend | 103023 | 1:200 | FC |
| CD45 Alexa Fluor 700 | Biolegend | 157210 | 1:200 | FC |
| CD45 Brilliant Violet 510 | Biolegend | 157219 | 1:200 | FC |
| CD45 Brilliant Violet 605 | Biolegend | 103140 | 1:200 | FC |
| CD45 PE | Biolegend | 103106 | 1:200 | FC |
| CD45 purified | BD | 550539 | 1:100 | IF |
| CD62L Brilliant Violet 421 | Biolegend | 104435 | 1:100 | FC |
| CD8 Alexa Fluor 488 | Biolegend | 100726 | 1:200 | FC |
| CD8 Brilliant Violet 650 | Biolegend | 100741 | 1:200 | FC |
| CD8 FITC | Biolegend | 100705 | 1:200 | FC |
| CD8 PE/Cy7 | Biolegend | 100721 | 1:200 | FC |
| CD8 purified | abcam | Ab22378 | 1:200 | IF |
| CD8a | BioXcell | BE0117 | 5-10 mg/kg | *in vivo* |
| CTLA-4 | BioXcell | BP0131 | 5 mg/kg | *in vivo* |
| F4/80 Alexa Fluor 647 | Serotec | MCA497A647 | 1:100 | IF |
| F4/80 PerCP/C5.5 | Biolegend | 123127 | 1:100 | FC, IF |
| gp100 (PMEL) PE | abcam | AB246731 | 1:200 | FC, IF |
| Ki67 Brilliant Violet 650 | Biolegend | 151215 | 1:200 | FC |
| KLRG1 Brilliant Violet 711 | BD | 564014 | 1:200 | FC |
| Lyve-1 | abcam | ab14917 | 1:100 | IF |
| MHC-I:SIINFEKL APC | Biolegend | 141605 | 1:100 | FC |
| MHC-I:SIINFEKL PE | Biolegend | 141603 | 1:100 | FC |
| MHC-II Alexa Fluor 647 | Biolegend | 115208 | 1:200 | FC |
| Pan-Cytokeratin Alexa Fluor 647 | Biolegend | 628604 | 1:100 | FC |
| Pan-Cytokeratin FITC | Sigma | F0397 | 1:100 | FC, IF |
| PD1 | BioXcell | BP0146 | 10 mg/kg | *in vivo* |
| PD1 APC-Fire 810 | Biolegend | 135251 | 1:200 | FC |
| PD1 Brilliant Violet 785 | Biolegend | 329929 | 1:200 | FC |
| Rat IgG controls | Jackson ImmunoResearch | 012-000-003 | 5-10 mg/kg | *in vivo* |
| Syrian Hamster | Leinco Technologies | I-444 | 5 mg/kg | *in vivo* |
| T-bet Pacific Blue | Biolegend | 644807 | 1:200 | FC |
| TCF1 FITC | BD | 567018 | 1:200 | FC |
| TCF1 PE | Biolegend | 655207 | 1:200 | FC |
| TCR-beta Brilliant Violet 711 | Biolegend | 109243 | 1:200 | FC |
| Tetramer (SIINFEKL) APC | Tetramer Core NIH | | 1:100 | FC |
| Tim3 Brilliant Violet 750 | BD | 755164 | 1:200 | FC |
| XCR1 APC/Cy7 | Biolegend | 148223 | 1:200 | FC |
| XCR1 Spark-UV-387 | Biolegend | 127677 | 1:200 | FC |

| Chemicals, Reagents, Kits | Source | Cat# | Concentration |
|---|---|---|---|
| Collagenase-IV | Sigma | C4-28 | 150 units/mL |
| DNase1 | Roche | 10104159001 | 100 µg/mL |
| FTY720 | MedChemExpress | HY-12005 | 2.5 µg/mL |
| TRITC-dextran 4.4 kDa | Sigma | T1037-50MG | 50 mg/mL |
| FITC-dextran 2,000 kDa | Sigma | FD2000S | 25 mg/mL |
| Vectashield Antifade Mounting Medium | Vector Laboratories | X-93952-24 | |
| DAPI | Sigma | D9542 | |
| Zombie UV | Biolegend | 423108 | 1:200 |
| GM-CSF | PeproTech | 315-03 | 20 ng/mL |
| IL-4 | PeproTech | 214-14 | 10 ng/mL |
| Pen/Strep | Gibco | 15140122 | 1 % |
| Sucrose | Sigma | S0389 | 30 % |
| β-Mercaptoethanol | Sigma | M3148 | |
| OCT compound | Tissue-Tek | FIS 23730571 | |
| LPS | Sigma | L-2630-10MG | 2 µg/mL |
| $OVA_{257-264}$ | Invitrogen | Vac-sin | |
| Normal donkey serum | Vector Laboratories | S-1000-20 | 5 % in PBS |
| Transcription Factor Fix/Perm Kit | eBioscience | 00-5521-00 | |
| CytoFix | BD | 554655 | |
| Fugene | Promega | E2311 | |
| PCR Mycoplasma Test Kit I/C | Promokine | PK-CA91-102 | |
| Fetal bovine serum | GeminiBio | 100-106 | 10% |
| Non-essential amino acids | Gibco | 11140050 | 1% |

**Image processing**

All image processing was done in ImageJ (https://imagej.nih.gov/ij/) or QuPath (https://qupath.github.io) to adjust brightness and contrast. Constant settings were used for all images from individual experiments. For signal quantification, Cellprofiler (https://cellprofiler.org) and Qupath were used. Tissue borders were identified manually, fluorescence signal intensity or area was identified by adaptive thresholding-minimum cross-entropy-algorithm. Positive signal objects were de-clumped by intensity (Cellprofiler), or positive cell detection command with manual threshold (QuPath). Object size-based filters were applied during cell identification (1-10 µm) to remove non-specific signal.

**Mathematical modeling**

The following steps were used to develop the numerical model presented in the main manuscript:

1. Create a deep network of lymph vessels and lymph nodes
2. Create a network of skin lymphatic vessels
3. Connect the deep and skin lymphatic vessel networks
4. Formulate equations to calculate pressure and flow through the networks
5. Perform dimensional analysis and derive analytical solution
6. Validate the pressure calculation with analytical and simplified numerical calculations
7. Make parameter estimates from the literature and simplified analyses
8. Perform a parametric study to determine sensitivities

***Deep network creation***

The network of deep lymphatic vessels and nodes for the human model was based on Savinkov *et al.*[1]. They generated the network from data obtained from PlasticBoy [Plasticboy. Plasticboy Pictures 2009 CC. http://www.plasticboy.co.uk/store/Human_Lymphatic_System_no_textures.html, (accessed on 21 December 2017)]. Their supplemental data provide two files. The first has the 'graphedges' which has the adjacency information with columns: 'From', 'To' and 'Length' where 'From' and 'To' are vertex numbers and 'Length' is the vessel length in mm. The second file is 'graphvertices' with columns: 'X', 'Y', 'Z', and 'isLymphNode', where the coordinates are in mm and 'isLymphNode' is 0 (not a node) or 1 (is a node). There are 996 vertices of which 272 are lymph nodes, 357 are the entrances to collecting vessels and the remainder are intermediate points along the collecting vessels. The same deep network was employed for the male and female human models.

A network with 43 lymph nodes in the mouse was based on the topology from Grebennikov *et al.*[2] with some modifications in the inguinal region and internal organs based on Kawashima *et al.*[3], Van den Broek *et al.*[4], and Harrell *et al.*[5]. Grebennikov *et al.*[2] did not include spatial coordinates, only the network connectivity. The spatial locations of the collecting vertices were manually adjusted to closely reproduce the drainage basins reported by Suami *et al.*[6] in the rat. The spatial coordinates of the lymph nodes themselves do not influence our results, only the connectivity of the deep network and the coordinates of the collecting vessels.

We acknowledge that considerable variability in the lymphatic networks may exist among human individuals, particularly in some regions of the human anatomy such as the head and neck[7–10]**.** The lymphatic network in mice also shows variability, albeit somewhat less than seen in humans[4]**.** As presented, our current maps may be best interpreted as being typical cases rather than attempts to perfectly model a specific patient or mouse. Nonetheless, the numerical methods presented here are sufficiently general to be readily adapted to alternative maps of the lymphatic system as they become available.

***Skin network creation***

The 3-D skin networks of the man and woman (**Fig. SM1**) were created from an .obj file of the integumentary system purchased from PlasticBoy [Plasticboy. Plasticboy Pictures 2009 CC. http://www.plasticboy.co.uk/store/Human_Male_Body_Textured_V04.html; http://www.plasticboy.co.uk/store/Human_Female_Body_Textured_V04.html, (accessed on 22 February 2023)] based on the same model as used for the deep lymphatic system in Savinkov *et al.*[1]. The .obj file was imported into Meshmixer where it was converted to a binary. stl format after remeshing to provide a finer and more uniform mesh. A typical mesh had more than 4,000 vertices. Minor corrections and smoothing were performed manually. The .stl file was read into Matlab with the utility stlread obtained from the Mathworks users group. We then converted the raw data from the .stl file from vertex and facet form to vertices, adjacency, flow conductance, area, and degree. The 3-D skin network of the mouse was created from an .stl file drawn in-house with a mesh created by a similar process in Meshmixer that was then converted to vertex, adjacency, flow conductance, area, and degree form.

It is important to note that the skin network in these models is a computational mesh used to calculate the flow. Resolutions of several centimeters and several millimeters are adequate for the human and mouse networks, respectively. The actual network of dermal lymphatic capillaries is much finer. The computational mesh needs to only represent the general 3D contours of the skin and the boundaries of the various lymphosomes.

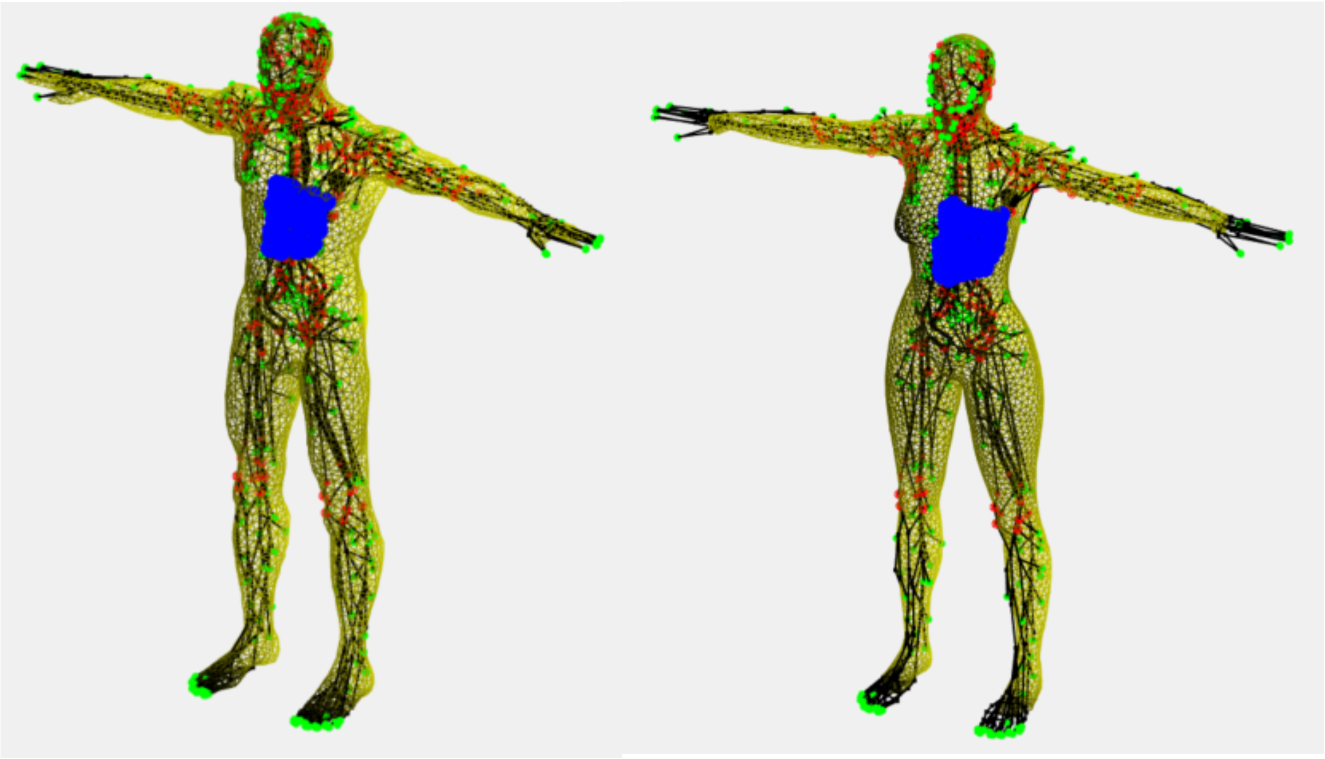

**Figure SM1. Male and female human models showing the drainage basin (blue) for a left axillary lymph node (black).**

***Connecting deep and skin networks***

A Matlab script was used to identify which of the deep vertices are collectors for the skin and which were collectors for deep organs. Then the script finds the nearest skin collector for each skin vertex. Additional scripts were used to identify the sentinel (first downstream) lymph node for each collector vertex. These tools were used to define the drainage basins (lymphosomes) associated with each node.

***Governing equations***

Flow follows interstitial pressure gradients (IFP) $p_i$ according to Darcy's law

$$\vec{u} = -\frac{\kappa}{\mu} \nabla p_i \tag{1}$$

where $\vec{u}$ is the velocity of fluid through the tissue, $\kappa$ is the tissue permeability and $\mu$ is the fluid viscosity (the ratio $\kappa/\mu$ is the hydraulic conductivity). The permeability includes fluid pathways through the interstitial space between cells as well as flow in the network of initial lymphatics that allow free fluid movement along pressure gradients due to their lack of intraluminal valves and contractility. The overall movement of fluid satisfies a mass balance given by equation below[11]:

$$\underbrace{\frac{\kappa}{\mu}\nabla\cdot\vec{u}}_{\substack{Lateral\\ Dermal\\ Flow}} = \underbrace{\beta_c(p_c - p_i)}_{\substack{Capillary\\ Leakage}} + \underbrace{\beta_L(p_L - p_i)}_{\substack{Lymphatic\\ Absorption}} \tag{2}$$

where the first and second terms on the righthand side are the distributed capillary and lymphatic sources or sinks of fluid, respectively (source when positive, sink when negative). Starling's law is used to represent the capillary source where the capillary conductance $\beta_c = L_p S_v$ with $L_p$ the capillary permeability and $S_v$ the surface area per unit volume. The capillary pressure can include osmotic effects $p_c = p_0 - \sigma\Delta\pi$ where $p_0$ is the mechanical pressure, $\sigma$ is the osmotic reflection coefficient and $\Delta\pi$ is the osmotic pressure difference) and is assigned at the upper boundary. A similar, simplified model of lymphatic clearance is used for the lymphatic clearance where $\beta_L$ is an effective lymphatic conductance with lymphatic pumping normally able to create a net suction effect ($p_L < p_i$), which if reduced has been demonstrated to contribute to edema. While more sophisticated models of lymphatic pumping are available, this two-parameter ($\beta_L, p_L$) model allows us to explore loss of lymphatic function by reducing $\beta_L$ or by making $p_L$ less negative to yield less suction or even positive to apply a retrograde pressure. Local equilibrium ($\vec{u} = 0$) near atmospheric pressure ($p_i = 0$) requires that $L_p S_v p_c = -\beta_L p_L$. When local lymphatic clearance is suppressed, fluid may be forced to adjacent lymphosomes ($\vec{u} \neq 0$) by small pressure gradients.

Interstitial pressures and flows were calculated by replacing the continuum formulation for the skin with a network of discrete tubes that represented ensembles of lymph capillaries and interstitial channels. Each tube provided the appropriate conductance for flow between points on the skin much like a finite difference approximation to the differential equations. In order to represent the continuum equations with the discrete vessels in the network model, we introduced the following mass balance analogous to that for the continuum equation

$$\underbrace{\sum_{j=1}^{degree_i} G_{skin}(p_j - p_i)}_{Lateral\ Dermal\ Flow} + \underbrace{G_c(p_c - p_i)}_{\substack{Capillary\\ Leakage}} + \underbrace{G_L(p_L - p_i)}_{\substack{Lymphatic\\ Absorption}} = 0 \tag{3}$$

where each flow was obtained from $G(p_j - p_i)$ and where the conductances were $G_{skin} = \frac{t\kappa}{\mu n}$—where *t* was the thickness of the skin and *n* was the number of segments per unit segment length (typically about 1.7 for networks where degree-6 vertices predominate), $G_c = tA_i\beta_c$ and $G_L = tA_i\beta_L$ where $A_i$ are the areas of associated skin vertices. We note that the skin thickness cancels out of each term. Here, the area associated with a vertex is approximated as a circle with a diameter based on the mean of the distances $d_{ij}$ to the $n$ connected vertices $j$

$$A_i \approx \frac{\pi}{4}\left\{\frac{1}{n}\sum_{j=1}^{n} d_{ij}\right\}^2 \tag{4}$$

At equilibrium, when all lymphosomes were enabled, we expected that little lateral movement of fluid occurred and that the interstitial pressure was near atmospheric so that $G_c p_c = -G_L p_L$. When a lymphosome was disabled, we set $G_L = 0$ in that region.

The resulting system of simultaneous equations was solved with Matlab for the interstitial pressure at each vertex and then for the flow between vertices.

***Dimensional analysis and analytical solution***

A planar, one-dimensional analysis provided validation and insight into the results from the full model. We considered a domain where $0 < x < R$ had the lymphatic collection disabled ($\beta_L = 0$) and $x > R$ where lymph clearance was enabled with $\beta_L > 0$. We assumed no flow at $x = 0$. The characteristic lengths over which the interstitial pressure decays exponentially were

$$L_{enabled} = \left(\frac{\kappa}{\mu(\beta_c + \beta_L)}\right)^{\frac{1}{2}} \tag{5}$$

and

$$L_{disabled} = \left(\frac{\kappa}{\mu\beta_c}\right)^{\frac{1}{2}} \tag{6}$$

The pressure in the disabled region ($0 < x < R$) is

$$p_i = p_c + C_1 \cosh\left(\frac{x}{L_{disabled}}\right) \tag{7}$$

and for the enabled region ($x > R$)

$$p_i = p_* + C_2 \exp\left(-\frac{x}{L_{enabled}}\right) \tag{8}$$

where $p_* = (\beta_c p_c + \beta_L p_L)/(\beta_c + \beta_L)$ and the constants are solutions of

$$\begin{bmatrix} cosh\left(\frac{R}{L_{disabled}}\right) & -\exp\left(-\frac{R}{L_{enabled}}\right) \\ \frac{1}{L_{disabled}} sinh\left(\frac{R}{L_{disabled}}\right) & \frac{1}{L_{enabled}} \exp\left(-\frac{R}{L_{enabled}}\right) \end{bmatrix} \begin{bmatrix} C_1 \\ C_2 \end{bmatrix} = \begin{bmatrix} p_* - p_c \\ 0 \end{bmatrix} \tag{9}$$

An analogous model using a planar, circular lymphosome can be obtained in terms of modified Bessel's functions ($I_0$, $K_0$, $I_1$, $K_1$) for the disabled region ($0 < r < R$)

$$p_i = p_c + C_1 I_0\left(\frac{r}{L_{disabled}}\right) \tag{10}$$

and for the enabled region ($r > R$)

$$p_i = p_* + C_2 K_0\left(\frac{r}{L_{enabled}}\right) \tag{11}$$

where the constants are solutions of

$$\begin{bmatrix} I_0\left(\frac{R}{L_{disabled}}\right) & -K_0\left(\frac{R}{L_{enabled}}\right) \\ \frac{1}{L_{disabled}} I_1\left(\frac{R}{L_{disabled}}\right) & \frac{1}{L_{enabled}} K_1\left(\frac{R}{L_{enabled}}\right) \end{bmatrix} \begin{bmatrix} C_1 \\ C_2 \end{bmatrix} = \begin{bmatrix} p_* - p_c \\ 0 \end{bmatrix} \tag{11}$$

The fluid velocity at the perimeter of the disabled region is given by

$$u = \frac{\kappa C_2}{\mu L_{enabled}} K_1\left(\frac{R}{L_{enabled}}\right) \tag{12}$$

The total fluid lost from the perimeter of the disabled region is given by

$$Q = \frac{2\pi R t \kappa C_2}{\mu L_{enabled}} K_1 \left(\frac{R}{L_{enabled}}\right) \tag{13}$$

***Validation***

Validation of the numerical model was performed by comparison with the analytical solutions. First, a planar, 1D model was considered where the analytical solution above was compared to the numerical solution that was obtained with a uniform mesh (**Figure SM2**) using the baseline parameters provided in **Table SM2** below.

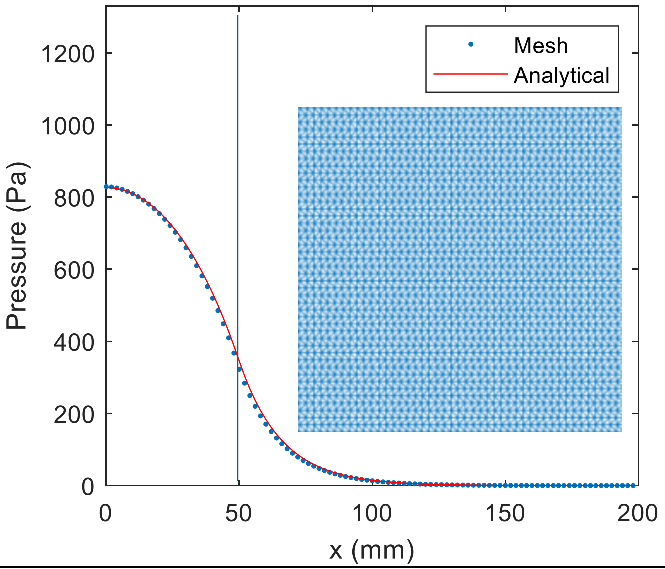


**Figure SM2. Analytical and uniform mesh results compared.** Vertical blue line marks the edge of the disabled lymphosome. A sample of the uniform mesh is shown as an insert.

An additional comparison was performed for the disabled human axillary lymphosome in the full model shown in **Fig. 4I** with the analytical result for a circular lymphosome using the same parameters with the surface area matched to give a nominal radius of disabled lymphosome = 65.8 mm (**Figure S4D-F)**. The slight shift of distance seen in the full 3D model is due to the curvature of the skin surface and the slightly non-circular boundary.

#### ***Parameter estimation***

Baseline values for the physical parameters are available in Table SM2. A more detailed discussion of the parameters and the sensitivity of the results to their variation follows.

**Table SM2. Baseline parameter values**

| Parameter | Description | Value |
|---|---|---|
| $\mu$ | viscosity | $1 \times 10^{-3}$ *Pa·s* |
| $\kappa$ | skin permeability | $3 \times 10^{-14}$ $m^2$ |
| $\delta$ | skin thickness | 0.5 to 2 *mm* |
| $D$ | lymph capillary diameter | 20 to 70 *μm* |
| $s$ | lymph capillary spacing | 400 *μm* |
| $\beta_c = L_p S_v$ | blood capillary conductance | $2 \times 10^{-8}$ $(Pa \cdot s)^{-1}$ |
| $\beta_L$ | lymph capillary conductance | $1 \times 10^{-7}$ $(Pa \cdot s)^{-1}$ |
| $L_{enabled}$ | absorption length with lymph clearance | 15.8 *mm* |
| $L_{disabled}$ | absorption length without lymph clearance | 38.7 *mm* |
| $n$ | density of lymph segments | 1.7 |
| $p_c$ | blood capillary pressure | 1330 *Pa* |
| $p_L$ | lymph suction pressure | -266 *Pa* |
| $R$ | lymphosome radius | 50 *mm* |

The lymphatic vessel segments in the skin network form a computational mesh where each segment represented ensembles of the dermal initial lymphatic vessels that are in fact much smaller and far more numerous than the network we used to model lateral flow within the skin. That is, the computational network represented the overall permeability of the tissue and its associated vessels, not a one-to-one model of the actual microvasculature. Values of the tissue permeability in the skin were obtained from measurements in similar tissues. See for example, values from Swartz *et al.*[12] in **Table SM3**. These were presented as being the permeability of the interstitial space alone, but it is unclear whether they might also include flow in the valve-less superficial lymph capillaries. The value from edematous tissue is relevant here because we expect some degree of edema in tissues when a lymphosome is disabled.

**Table SM3. Tissue permeability values (κ in m²) from Swartz et al**[12]

| Method | Normal | Edema |
|---|---|---|
| Fluorescence | $2.43 \times 10^{-16}$ | $2.26 \times 10^{-14}$ |
| Pipet | $3.32 \times 10^{-14}$ | |

Alternatively, the skin permeability was estimated from the typical dimensions and spacing of the initial lymphatic vessels. We estimated the thickness $\delta$ of the skin to be 0.5 to 2 mm, the spacing $s$ of initial lymphatic vessels to be 400 μm, and their diameters D to be 20 to 70 μm. Assuming Poiseuille flow in each parallel capillary, the equivalent permeability was estimated from Truskey *et al.*[13].

$$\kappa = \frac{\pi D^4}{128 s \delta} \tag{14}$$

The range of permeabilities arising from the range of parameters given was $\kappa = 5x10^{-15}$ to $3x10^{-12}\ m^2$ which was similar to, but somewhat expanded relative to the range observed by Swartz *et al*. This estimate was highly sensitive to the vessel diameter, but as we show later, some important results are relatively insensitive to $\kappa$.

#### *Parametric sensitivity*

We performed a parametric sensitivity study using the analytic solution for a disabled, circular lymphosome surrounded by normal tissue (Eqns. 11-13) (**Table SM4**). Of particular interest is the interstitial pressure at the center of the disabled lymphosome, which can approach the local capillary blood pressure as a limit, and the quantity and destination of fluid diverted from the disabled lymphosome to adjacent absorption sites. These quantities depend on the radius of the lymph disabled lymphosome $R$, the capillary blood pressure $p_c$, lymph suction pressure $p_L$, blood capillary conductance $\beta_c = L_p S_v$, skin permeability $\kappa$ , and lymph vessel conductance $\beta_L$. While we provide baseline estimates of each parameter, considerable physiological variability can be expected in each parameter. Moreover, the parameters can be significantly modified by physiological conditions such as fitness, widespread inflammation and obesity with or without inflammation[14–18]. Figure S4A-C presents simulated results showing how doubling or halving some of these parameters can change measures of clinical relevance for a disabled axillary lymphosome surrounded by functional lymph clearance. We expect obesity to reduce tissue permeability and lymph pumping capacity leading to increases in interstitial pressures and a reduced rate of antigen carrying fluid to adjacent lymphosomes. Whereas physical fitness is predicted to compensate for these effects. Widespread inflammation, which often accompanies obesity, tends to increase capillary permeability leading to increased interstitial pressure and rate of fluid diversion. While the total quantity of diverted fluid varies among the conditions considered, the model predicts little difference in how the diverted fluid is apportioned among the neighboring lymphosomes.

Most of the results depend directly on $R/L_{enabled}$ and $R/L_{disabled}$, the ratios of the lymphosome radius to the characteristic lengths over which interstitial pressures vary (Equations 5 and 6). Typical values of $L_{enabled}$ and $L_{disabled}$ fall in the range of tens of millimeters. As explained above, theoretical estimates and measurements of the tissue permeability vary widely. Fortunately, since the characteristic lengths have a square root dependence on the tissue permeability, the characteristic lengths are relatively insensitive to tissue permeability. For example, a 100-fold change in tissue permeability changes the lengths by only a factor of 10.

Changes in the characteristic lengths $R/L$ do affect how closely the interstitial pressure in the center of a disabled lymphosome tracks the blood pressure in the capillaries (**Figure S4D**). Tissues with large disabled lymphosomes that have low tissue permeability or that have highly permeable vessels are expected to have interstitial pressures that approach the capillary pressure leading to significant edema—a condition difficult to achieve in a small animal such as a mouse, but more likely to occur on a human scale. The total fluid diverted from a disabled lymphosome (Q in Equation 13) roughly increases with the area of the disabled lymphosome as expected due to the greater tissue volume that must find an alternative outlet for its interstitial fluid. Increases in the conductance of the blood capillaries ($\beta_c$) also increase the total fluid diverted, but with less than a proportional rate. The conductance of the lymph in the tissue surrounding the disabled lymphosome ($\beta_L$) has only a weak effect on the pressures and flow (**Figure S4D**). For example, increasing $\kappa$ by a factor of 10,000 resulted in an increase in the velocity of interstitial flow at the perimeter of the disabled lymphosome by only a factor of about 15 because changes in pressure gradient were offset by changes in $\kappa$.

The ultimate destination of fluid diverted from a disabled lymphosome has little dependence on the characteristic lengths. In nearly all cases, the displaced fluid will be absorbed by the lymphatic vessels in a lymphosome immediately adjacent to the disabled lymphosome. Only in exceptional cases, where the adjacent lymphosome is significantly smaller than the characteristic length, would fluid reach beyond the nearest neighbor. Nonetheless, fluid diverted from a disabled lymphosome may still reach a distant lymph node due to the routing of collecting lymphatic vessels from adjoining drainage basins.

The anatomical differences between mouse and human may alter the destination of the fluid. For example, the direct connection from the inguinal region to the axillary region that exists in the mouse is generally absent in humans. Disabling an axLN in a human did not affect drainage to the inguinal basin, but, like the mouse, diverted fluid to contralateral LNs (**Fig. 4I**). Moreover, disabling a single axLN in a human did not produce the same extensive re-routing of flow seen in the mouse because the human has numerous nodes in the axillary region that form a redundant network. Note that anatomical variations in the network can exist between individuals, and this will affect the flow redundancy.

**Table SM4. Parametric variation relative to baseline values (subscript 0) in Table SM3.** Note that negative interstitial pressures arise only if the lymph pumping is assumed to remain constant when parameters are changed.

| | $L_{disabled}$ $(mm)$ | $L_{enabled}$ $(mm)$ | $R/L_{disabled}$ | $R/L_{enabled}$ | $p_i(0)/p_c$ | $Q$ $(\frac{ml}{hr})$ |
|---|---|---|---|---|---|---|
| Baseline | 38.7 | 15.8 | 1.29 | 3.16 | 0.43 | 0.53 |
| $R = R_0 x 0.1$ | 38.7 | 15.8 | 0.129 | 0.316 | 0.16 | 0.03 |
| $R = R_0 x 10$ | 38.7 | 15.8 | 12.9 | 31.6 | 1.00 | 8.07 |
| $\kappa = \kappa_0 x 0.1$ | 12.2 | 5.0 | 4.10 | 10.0 | 0.94 | 0.24 |
| $\kappa = \kappa_0 x 10$ | 122 | 50.0 | 0.41 | 1.00 | 0.09 | 0.70 |
| $\beta_c = \beta_{c0} x 0.1$ | 122 | 17.1 | 0.41 | 2.92 | -0.10 | 0.08 |
| $\beta_c = \beta_{c0} x 10$ | 12.2 | 10.0 | 4.10 | 5.00 | 0.98 | 0.78 |
| $\beta_L = \beta_{L0} x 0.1$ | 38.7 | 31.6 | 1.29 | 1.58 | 0.80 | 0.19 |
| $\beta_L = \beta_{L0} x 10$ | 38.7 | 5.4 | 1.29 | 9.26 | 025 | 0.69 |

### Statistics

Statistical significance was determined using Mann-Whitney U test, two-way ANOVA test or Log-rank (Mantel-Cox) test. Data are presented as mean ± SEM. Fisher's exact test was used to analyze contingency tables including responders vs. non-responders and survivors vs. non-survivors. Statistical significance threshold was set at $p<0.05$. All calculations were performed using GraphPad Prism 9 software.
